# Growth-dependent signals drive an increase in early G1 cyclin concentration to link cell cycle entry with cell growth

**DOI:** 10.1101/2020.09.30.321182

**Authors:** Robert A. Sommer, Jerry T. DeWitt, Raymond Tan, Douglas R. Kellogg

**Affiliations:** Department of Molecular, Cell, and Developmental Biology, University of California, Santa Cruz, Santa Cruz, CA 95064, USA

## Abstract

Entry into the cell cycle occurs only when sufficient growth has occurred. In budding yeast, the cyclin Cln3 initiates cell cycle entry by inactivating a transcriptional repressor called Whi5. Growth-dependent changes in the concentrations of Cln3 or Whi5 have been proposed to link cell cycle entry to cell growth. However, there are conflicting reports regarding the behavior and roles of Cln3 and Whi5. Here, we found no evidence that changes in the concentration of Whi5 play a major role in controlling cell cycle entry. Rather, the data suggest that cell growth triggers cell cycle entry by driving an increase in the concentration of Cln3. We further found that accumulation of Cln3 is dependent upon homologs of mammalian SGK kinases that play roles in control of cell growth and size. Together, the data are consistent with models in which Cln3 serves as the crucial link between the cell cycle and signals that control cell growth and size.

## Introduction

Cell cycle progression is subservient to cell growth. Thus, key cell cycle transitions occur only when sufficient growth has occurred. A particularly important point at which cell growth controls cell cycle progression is at the end of G1 phase, when cells decide whether to commit to cell cycle entry. The key molecular event that marks cell cycle entry is transcription of late G1 phase cyclins, which bind and activate cyclin-dependent kinases to initiate cell cycle events. Transcription of the late G1 cyclins is initiated only when sufficient growth has occurred.

A key question concerns how growth triggers cell cycle entry. Most models suggest that growth-dependent changes in the number or concentration of key cell cycle regulatory proteins triggers cell cycle entry. If the number of molecules of a protein increases at the same rate as growth, the number of molecules per cell will increase but the concentration of the protein will stay constant. In this case, an increase in the number of protein molecules per cell could be sufficient to trigger cell cycle entry if the activity of the protein is titrated against an activity that does not scale with growth. Alternatively, cell cycle entry could require changes in the concentration of cell cycle regulatory proteins. In this case, if the number of molecules per cell increases faster than growth, the concentration and overall activity of the protein should also increase. Conversely, if the number of protein molecules stays constant as growth occurs the concentration of the molecule will decrease. It remains unclear whether changes in number or concentration of cell cycle regulators triggers cell cycle entry. In addition, it remains possible that changes in the catalytic rate of an enzyme, such as a kinase or phosphatase, play an important role.

In budding yeast, key regulators of the cell cycle that are thought to link cell cycle entry to cell growth were first identified via genetic analysis. An early G1 cyclin called Cln3 was found to be a dose-dependent regulator of cell cycle entry and cell size (Cross 1988; Nash *et al.* 1988). Overexpression of *CLN3* causes premature cell cycle entry at a reduced cell size, while loss of Cln3 causes delayed cell cycle entry at a large cell size. A poorly understood protein called Bck2 plays redundant roles with Cln3 (Epstein and Cross 1994). Loss of Cln3 or Bck2 causes delayed cell cycle entry, whereas loss of both causes a failure in cell cycle entry. The Cln3/Cdk1 complex triggers cell cycle entry by activating the SBF transcription factor, which drives transcription of late G1 phase cyclins as well as hundreds of additional genes required for subsequent cell cycle events. The Cln3/Cdk1 complex is thought to activate SBF by directly phosphorylating and inactivating Whi5, a transcriptional repressor that keeps SBF inactive prior to cell cycle entry (Jorgensen *et al.* 2002; Costanzo *et al.* 2004; de Bruin *et al.* 2004). Loss of Whi5 causes premature cell cycle entry at a reduced cell size, which suggests that Whi5 plays an important role in the mechanisms that link cell cycle entry to cell growth (Jorgensen *et al.* 2002).

Early studies suggested a model in which accumulation of Cln3 protein drives cell cycle entry (reviewed in Jorgensen and Tyers 2004b; Turner *et al.* 2012). Cln3 protein is rapidly turned over, which should make Cln3 levels highly sensitive to translation rate (Tyers *et al.* 1992). If translation rate increases with cell size, the number of Cln3 protein molecules in the cell should also increase with cell size, which could lead to cell cycle entry. Difficulties in detecting Cln3 protein initially made it difficult to test the prediction that Cln3 protein levels increase with cell growth during G1 phase. More recent studies found that levels of Cln3 increase gradually during G1 phase, reaching a peak around the time of bud emergence, consistent with the idea that gradually rising levels of Cln3 trigger cell cycle entry (Zapata *et al.* 2014). Another recent study used an in vivo reporter to measure the rate of Cln3 production and found that the rate increases faster than the rate of growth in G1 phase, which suggests that an increase in the concentration of Cln3 could trigger cell cycle entry (Litsios *et al.* 2019). In contrast, another study concluded that the concentration of Cln3 does not change during growth (Schmoller *et al.* 2015). Overall, the relationship between Cln3 accumulation and cell growth remains poorly understood. It is unclear whether Cln3 provides a simple readout of overall translation rate or whether the rate of Cln3 accumulation is influenced by other growth-dependent signals. It is also unclear how regulation of Cln3 transcription, translation, and turnover influence accumulation of Cln3 during cell growth. Finally, it remains unknown whether or how a threshold level of Cln3 triggers cell cycle entry.

Another study suggested that dilution of Whi5 by cell growth is the critical event that links cell cycle entry to cell growth (Schmoller *et al.* 2015). In this study, it was found that the concentration of Cln3 remains constant during growth because the number of Cln3 molecules per cell increases in a manner that is commensurate with growth. In contrast, the concentration of Whi5 was found to decrease because no new Whi5 is produced during G1 phase. The model postulates that once Whi5 concentration drops below a threshold, the activity of Cln3 becomes sufficient to inactivate Whi5, thereby triggering cell cycle entry. A limitation of this model is that cells undergo little growth in G1 phase (Ferrezuelo *et al.* 2012; Leitao and Kellogg 2017; Litsios *et al.* 2019), which means that Whi5 concentration could be reduced by less than 50%. It is unclear how such small changes in Whi5 concentration could be translated into an effective cell size control decision. Another limitation is that the effects of Whi5 protein concentration on cell cycle entry were analyzed only in *bck2Δ* cells, and only under nutrient poor conditions. Finally, the behavior of Cln3 protein was analyzed using a mutant version of Cln3 that lacks amino acid sequences that target it for proteolytic destruction, which results in overexpression and loss of cell cycle-dependent changes in protein levels.

Another limitation of current models for size control in G1 phase is that they fail to explain proportional relationships between cell size and growth rate that hold across all orders of life. Yeast cells growing slowly in poor nutrients are nearly 2-fold smaller than cells growing rapidly in rich nutrients (Johnston *et al.* 1977). Moreover, a proportional relationship between cell size and growth rate holds even when comparing yeast cells growing at different rates under identical nutrient conditions (Ferrezuelo *et al.* 2012; Leitao and Kellogg 2017). For example, cells in a population of wild type yeast cells in early G1 phase show a 3-fold variance in growth rate. The slow growing cells within the population undergo cell cycle entry at a reduced cell size compared to rapidly growing cells. Thus, it appears that signals linked to nutrients and growth rate modulate the threshold amount of growth required for cell cycle progression. Classic experiments in fission yeast suggest that the threshold amount of growth required for cell cycle progression is readjusted within minutes of a shift to new nutrient conditions (Fantes and Nurse 1977). A model that could explain the link between cell size and growth rate is that global signals that set growth rate also set the threshold amount of growth required for cell cycle progression (Lucena *et al.* 2018). However, the mechanisms by which nutrient-dependent signals modulate growth thresholds remain largely unknown.

Distinguishing models will require a clear understanding of how changing concentrations of Cln3 and Whi5 influence cell cycle entry, and how the concentrations of Cln3 and Whi5 vary during growth and in response to changes in growth rate. However, there are conflicting reports on the behaviors of Cln3 and Whi5 during growth in G1 phase (Tyers *et al.* 1993; Liu *et al.* 2015; Schmoller *et al.* 2015; Zapata *et al.* 2014; Lucena *et al.* 2018; Dorsey *et al.* 2018; Blank *et al.* 2018; Litsios *et al.* 2019; Barber *et al.* 2020). Most previous studies used quantitative fluorescence microscopy to analyze protein behavior in living cells. Differences in fluorescent tags, nutrient and imaging conditions, potential imaging artifacts, as well as the use of mutant proteins or mutant genetic backgrounds could explain differences in results. In addition, the low abundance of Cln3 has made it difficult to measure levels of wild type Cln3 by fluorescence microscopy.

Here, we used quantitative western blotting in synchronized cells to carry out a comprehensive analysis of the function and regulation of Cln3 and Whi5 during growth in G1 phase. A simple model for nutrient modulation of cell size could be that poor nutrients cause a reduction in Whi5 protein levels, which would reduce the amount of growth required to dilute Whi5 below the threshold required for cell cycle entry. However, little is known about the relationship between Cln3 and Whi5 protein levels during growth in G1 phase in differing nutrient conditions. We therefore investigated how levels of the Whi5 and Cln3 proteins are modulated during growth in G1, and how nutrient availability modulates levels of both proteins. We also investigated the mechanisms that control accumulation of Cln3 during growth in G1 phase.

## Results

### Cells growing in poor carbon undergo cell cycle entry at a dramatically lower ratio of Cln3 to Whi5

Previous studies found that levels of Cln3 mRNA and protein are reduced in asynchronous cells growing under poor nutrient conditions (Gallego *et al.* 1997; Parviz and Heideman 1998; Hall *et al.* 1998; Blank *et al.* 2018). However, more recent work discovered that Cln3 is also synthesized in mitosis (Landry *et al.* 2012; Zapata *et al.* 2014). Therefore, analysis of Cln3 in asynchronous cells could not define the behavior of Cln3 during growth in G1 phase. Here, we used synchronized cells to characterize the dynamics of Cln3 and Whi5 proteins during growth in G1 phase in rich or poor carbon.

Cells were grown in a poor carbon source (YP media containing 2% glycerol and 2% ethanol) and centrifugal elutriation was used to isolate small newborn daughter cells in early G1 phase. The isolated cells were then resuspended in YP media containing rich carbon (2% glucose) or poor carbon to initiate growth in G1 phase. An advantage of this approach is that the cells growing in rich versus poor carbon were previously grown under identical conditions and each culture contains identical numbers of cells, which allows comparison of levels of Cln3 and Whi5 between conditions. Another advantage is that the cells shifted to rich carbon undergo a prolonged growth interval in G1 phase to reach the increased size of cells in rich carbon, which provides a longer interval for examining the behavior of Cln3 and Whi5. A disadvantage of this approach is that the cells shifted from poor to rich carbon may show effects that are associated with adjusting metabolic pathways to utilize a rich carbon source.

Median cell size was determined with a Coulter Channelyzer and plotted as a function of time (**Figure 1A**). Since bud emergence provides a marker of cell cycle entry, the percentage of cells with buds was plotted as a function of time (**Figure 1B**). Finally, a plot of the percentage of cells with buds versus cell size provided a measure of cell size at cell cycle entry (**Figure 1C**). As expected, the cells transferred to rich carbon grew faster and spent more time undergoing growth in G1 phase compared to cells in poor carbon. The cells in rich carbon also entered the cell cycle at a larger size than the cells in poor carbon. The cells that remained in poor carbon grew in size by only 24% before bud emergence (defined as the time at which 20% of cells were budded). In contrast, the cells shifted from poor to rich carbon increased in size by 75%.

**Figure 1:**
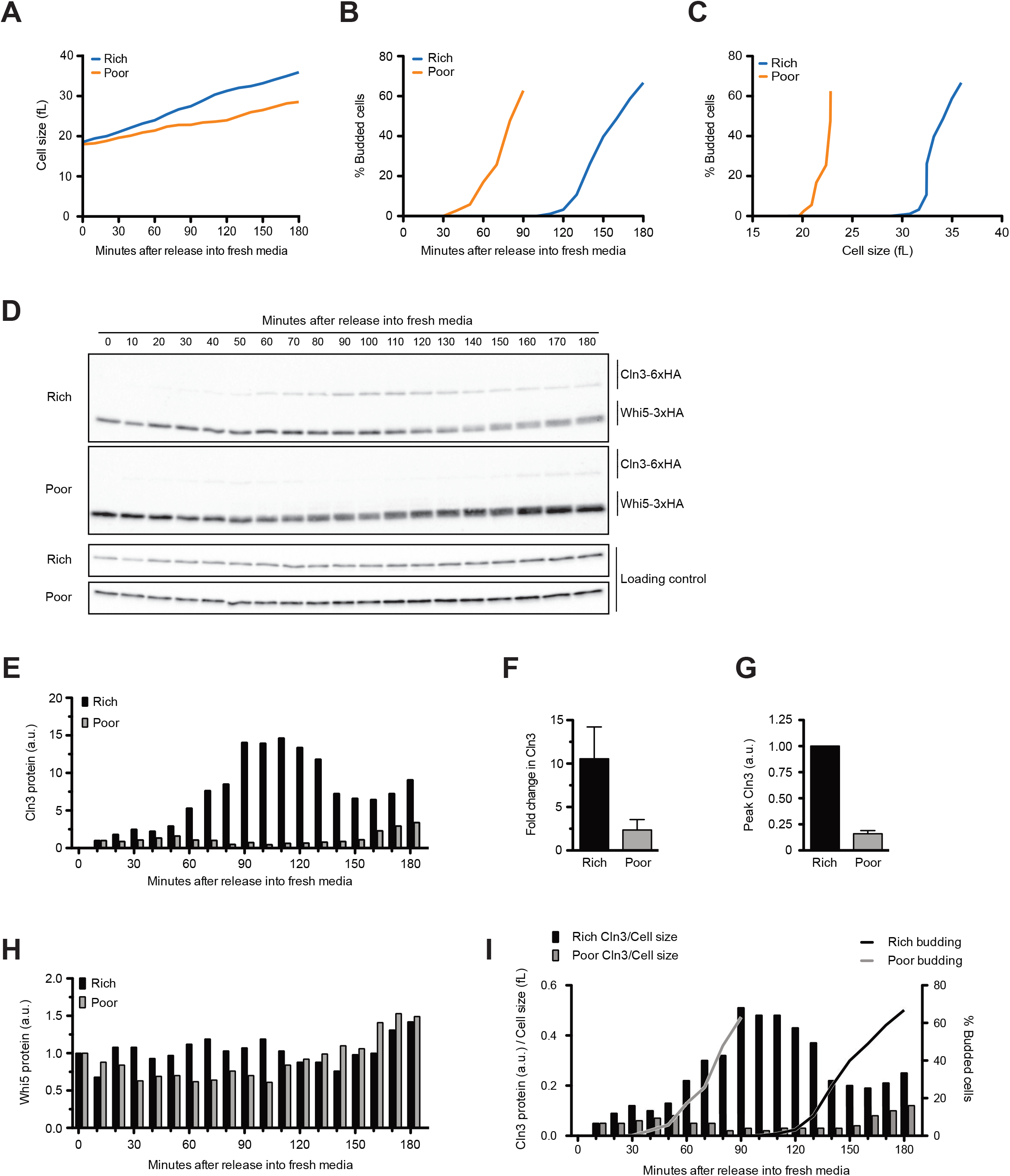
Wild type cells were grown to mid-log phase in poor carbon (YPG/E) and small unbudded cells were isolated by centrifugal elutriation. Cells were released in either rich carbon (YPD) or poor carbon (YPG/E) at 25°C and samples were taken at 10 minute intervals. All data in the Figure are from the same biological replicate. **(A)** Median cell size (fL) was measured using a Coulter Counter and plotted as a function of time. **(B)** The percentage of budded cells as a function of time. **(C)** The percentage of budded cells plotted as a function of median cell size at each time point. **(D)** The behavior of Cln3-6xHA and Whi5-3xHA were analyzed by western blot on the same blot for each carbon source. An anti-Nap1 antibody was used a loading control. **(E)** Quantification of Cln3-6xHA protein levels as a function of time. At each time point, the Cln3 signal was calculated as a ratio over the 10 min time point. **(F)** The fold change in Cln3 protein signal was calculated as a ratio of the signal from the time point with peak Cln3 over the signal from the 10 min time point. The data represent the average of three biological replicates, and the error bars represent SEM. **(G)** The difference in peak Cln3 between rich and poor carbon was calculated by first normalizing the peak Cln3 signal to the loading control in each carbon source, and then comparing the signal between carbon sources. **(H)** Quantification of Whi5-3xHA from western blots in (D). The change in protein abundance over time was measured by calculating the ratio over the zero time point for each time point. The Whi5 signal was not normalized to a loading control because the loading control signal increases gradually with growth. **(I)** The relative change in Cln3 concentration over time was calculated by taking the ratio of the Cln3 signal over cell size at each time point. The Cln3 signal was not normalized to a loading control because the loading control signal increases gradually with growth. The bud emergence data from panel (B) are included for reference. The experiment shown in this figure was repeated for 3 biological replicates, which gave similar results.

The behaviors of Whi5 and Cln3 were assayed in the same samples. Whi5 was detected with a 3XHA tag, whereas Cln3 was detected with a 6XHA tag. Cln3 tagged with 3XHA could not be detected, which necessitated use of the 6XHA tag. Tagging both Cln3 and Whi5 with 6XHA caused the proteins to co-migrate on SDS-PAGE, which prevented use of identical tags. The use of HA tags on both proteins allowed detection on the same western blot so that relative levels of Cln3 and Whi5 could be compared. However, the use of tags with different detection sensitivities precluded comparison of absolute protein levels between Cln3-6XHA and Whi5-3XHA. Nevertheless, the fact that Whi5-3XHA is easily detectable, whereas Cln3-3XHA is undetectable, indicates that levels of Whi5 far exceed those of Cln3. Western blots comparing protein levels in rich and poor carbon were carried out simultaneously under identical conditions to allow direct comparison of protein levels between carbon sources. Here, we will use “concentration” to refer to the number of molecules per unit of cellular volume, and the more general term “protein levels” to refer to the number of molecules in the cell.

In both conditions, Cln3 was not detectable at the zero time point (**Figure 1D, see Figure 1 – figure supplement 1A for a longer exposure**). Cln3 was first detected at 10 minutes after elutriation and levels of Cln3 increased gradually during growth in both rich and poor carbon. Cln3 levels peaked slightly before the first buds could be detected in both conditions. Cln3 levels were dramatically lower in poor carbon. Quantification of the Cln3 signal relative to the 10 minute time point in multiple biological replicates indicated that Cln3 protein levels increased approximately 10-fold before bud emergence in rich carbon and 2.5-fold in poor carbon (**Figures 1E and 1F)**. Quantification of peak Cln3 levels in rich and poor carbon suggest that cells in poor carbon initiate cell cycle entry with approximately 8-fold lower levels of Cln3 than cells in rich carbon (**Fig. 1G**).

We did not detect differences in Whi5 protein levels between the two carbon sources. In Figure 1D, a slightly longer exposure was used for the western blot from cells in poor carbon so that Cln3 could be detected, but quantification of Whi5 levels relative to a loading control showed no difference between rich and poor carbon. To test whether Whi5 levels change during G1 phase we calculated a ratio of the Whi5 signal at each time point over the Whi5 signal at the zero time point (**Figure 1H**). We used the same approach to calculate the fold change in Whi5 levels between the zero time point and bud emergence in multiple biological replicates (**Figure 1 – figure supplement 1B**). Both plots suggest that levels of the Whi5 protein do not change substantially before bud emergence in rich or poor carbon, consistent with several previous studies that used fluorescence microscopy to analyze Whi5 levels (Schmoller *et al.* 2015; Litsios *et al.* 2019).

A fraction of Whi5 shifted to lower electrophoretic mobility forms at the time of bud emergence, which likely corresponds to the activity of a previously described positive feedback loop in which the late G1 cyclins Cln1 and Cln2 activate Cdk1 to phosphorylate and inactivate Whi5 (Cross and Tinkelenberg 1991; Costanzo *et al.* 2004).

Together, the data show that Cln3 protein levels are correlated with the extent of growth in volume in G1 phase in both rich and poor carbon, reaching a peak near the time of bud emergence. In addition, overall Cln3 protein levels in G1 phase are correlated with the growth rate set by carbon source. Thus, Cln3 levels are high in rich carbon and low in poor carbon. In contrast, Whi5 protein levels are not modulated by carbon source. As a result, cells growing in poor carbon enter the cell cycle at a much lower ratio of Cln3 to Whi5 protein compared to cells in rich carbon. We estimate that there is a nearly 8-fold increase in the ratio of Whi5 to Cln3 at cell cycle entry in poor carbon compared to rich carbon. These results suggest that the threshold amount of Cln3 needed to trigger cell cycle entry is modulated by nutrient availability.

### The concentration of Cln3 increases during growth in G1 phase

A previous study utilized quantitative fluorescence microscopy to analyze accumulation of a mutant stabilized version of Cln3 and concluded that the concentration of Cln3 does not change during growth in poor carbon (Schmoller *et al.* 2015). To estimate changes in wildtype Cln3 concentration before bud emergence, we divided the Cln3 signal from western blots by cell volume at each time point (**Figure 1I**). Using this approach, we quantified the increase in Cln3 concentration between the 10 minute time point and the time of bud emergence for multiple biological replicates (**Figure 1 – figure supplement 1C**). This showed that the concentration of Cln3 increased 7-fold prior to bud emergence in rich carbon, and 2-fold in poor carbon. A rise in Cln3 concentration during G1 phase was also detected in a previous study that used targeted proteomics to analyze Cln3 concentration (Litsios *et al.* 2019).

We carried out a similar analysis for Whi5 and found that Whi5 concentration decreased by approximately 30% in rich carbon and 20% in poor carbon, consistent with the fact that Whi5 protein levels do not change substantially and there is only a small increase in cell size in G1 phase (**Figure 1 – figure supplement 1D**). A decrease in the concentration of Whi5 during G1 phase was also detected in a previous study (Schmoller *et al.* 2015), although the decrease in concentration observed here was substantially smaller. Our results are consistent with several previous studies that observed a relatively small increase in cell volume in G1 phase (Ferrezuelo *et al.* 2012; Leitao and Kellogg 2017) and little change in Whi5 concentration (Litsios *et al.* 2019).

### Cln3 protein levels respond rapidly to changes in nutrient availability

In both budding yeast and fission yeast, cells rapidly readjust the threshold amount of growth required for cell cycle progression when shifted to nutrient conditions that support different growth rates (Fantes and Nurse 1977; Johnston *et al.* 1979; Lucena *et al.* 2018). In fission yeast, the threshold appears to be readjusted within minutes. Therefore, cell cycle regulators that link cell cycle progression to cell growth should show a similar rapid response to changes in nutrients.

A previous study found that Cln3 protein levels decrease within 30 minutes of a shift from rich to poor nitrogen, but did not test shorter time points or the effects of carbon source (Gallego *et al.* 1997). We therefore analyzed the behavior of Cln3 and Whi5 following a shift from rich to poor carbon. Cln3 underwent rapid changes in abundance when asynchronous cells were shifted from rich to poor carbon (**Figure 2**). The amount of Cln3 protein transiently increased within 5 minutes and then rapidly disappeared so that it was nearly undetectable. In contrast, Whi5 levels remained constant (**Figure 2**). Whi5 phosphorylation was slowly lost during the time course, consistent with previous studies showing that cells in poor nutrients spend more time in early G1 phase, which is when Whi5 is found in a dephosphorylated state (Hartwell and Unger 1977). These observations show that Cln3, but not Whi5, responds rapidly to nutrient-dependent signals that modulate cell growth and size.

**Figure 2:**
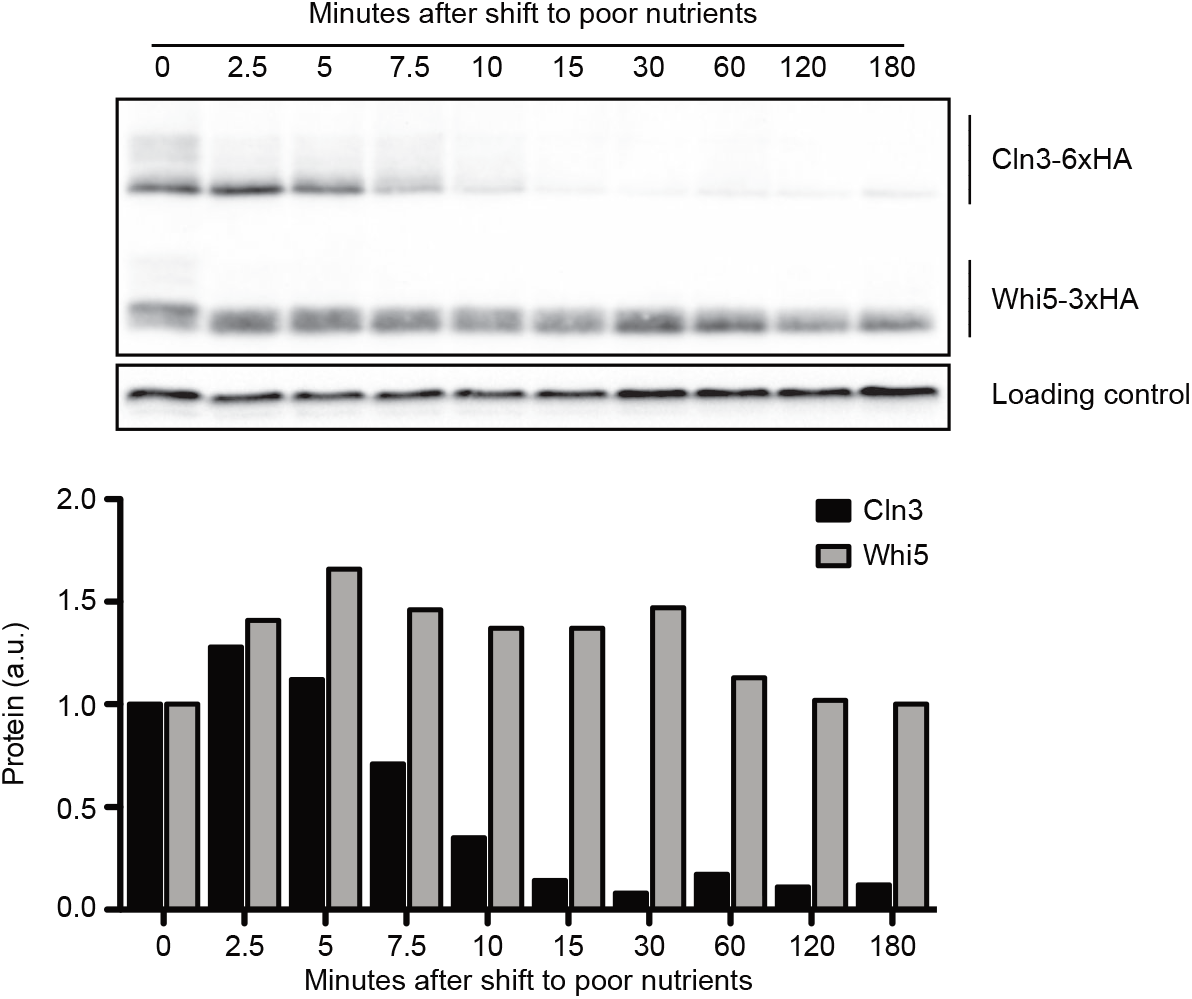
Rapidly growing cells in rich carbon (YPD) were shifted to poor carbon (YPG/E) at 25°C and the behavior of Cln3-6xHA and Whi5-3xHA was analyzed by western blot on the same blot. The levels of Cln3 and Whi5 protein were quantified relative to the 0 min time point. An anti-Nap1 antibody was used as a loading control.

### Overexpression of *WHI5* does not cause large effects on cell size

The discovery that cells in rich or poor carbon undergo cell cycle entry with a large difference in the ratio of Whi5 to Cln3 suggests that Whi5 protein concentration may not have a major influence on cell cycle entry. To investigate further, we carried out additional tests of how the concentration of Whi5 influences cell cycle entry and cell size. A previous study found that an additional copy of *WHI5* causes an increase in cell size (Schmoller *et al.* 2015). However, the effects of an additional copy of *WHI5* were only tested under nutrient-poor condition and in cells that lack Bck2, a poorly understood inducer of cell cycle entry that becomes essential in cells that lack Cln3. In addition, the extra copy of *WHI5* was tagged, which could influence function, and it included both the promoter and part of the coding sequence of a gene neighboring *WHI5*.

We constructed an integrating vector that includes only the wild type untagged *WHI5* gene with the normal upstream and downstream control regions. The *WHI5* construct fully rescued the size defects of *whi5Δ* (**Figure 3 – figure supplement 1A**). Integration of the plasmid into wild type cells to introduce an extra copy of *WHI5* had little effect on the size of cells growing in YP medium containing rich or poor carbon (**Figure 3A**). When cells were grown in nutrient-poor synthetic complete medium the extra copy of *WHI5* caused a 7% increase in median cell size in rich carbon and a 10% increase in poor carbon (**Figure 3 – figure supplement 1B**), similar to the previously reported effects of an extra copy of *WHI5* in *bck2Δ* cells growing in synthetic complete media (Schmoller *et al.* 2015). The finding that an extra copy of *WHI5* has stronger effects on cell size in nutrient poor synthetic complete media may be due to the large reduction in Cln3 protein levels in poor nutrients.

**Figure 3:**
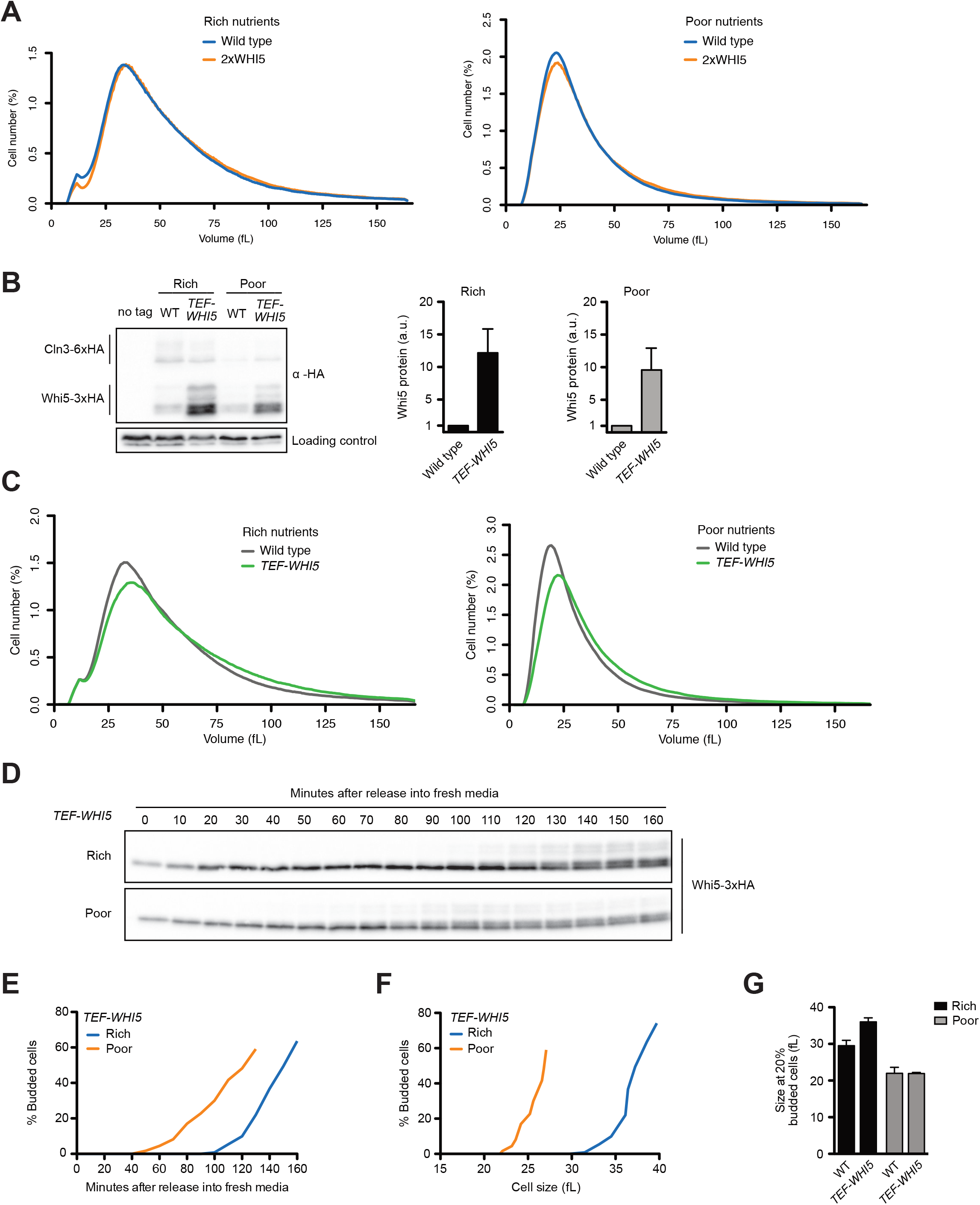
**(A)** Wild type and *2xWHI5* cells were grown to log phase in rich (YPD) or poor carbon (YPG/E) and cell size was measured using a Coulter Counter. **(B)** Cells that express *CLN3-6XHA* and either *WHI5-3XHA* or *TEF1-WHI5-3XHA* were grown to mid log phase in rich carbon (YPD) or poor carbon (YPG/E). Cln3-6xHA and Whi5-3xHA were imaged on the same blot. An anti-Nap1 antibody was used as a loading control. The *TEF1-WHI5-3XHA* signal was quantified relative to the WHI5-3XHA signal after each signal was normalized to the loading control. **(C)** Wild type and *TEF-WHI5* cells were grown to log phase in rich (YPD) or poor carbon (YPG/E) and cell size was measured using a Coulter Counter. **(D-F)** *TEF-WHI5* cells were grown to mid log phase in poor carbon (YPG/E) and small unbudded cells were isolated via centrifugal elutriation. Cells were then released into rich (YPD) or poor (YPG/E) carbon and samples were collected at 10 minute intervals. **(D)** The behavior of Whi5-3xHA was analyzed by western blot. See **Figure 3 – figure supplement 1E** for a loading control. **(E)** Percentage of budded cells as a marker for cell cycle entry. **(F)** Median cell size was measured using a Coulter counter and plotted against bud emergence to show cell size at cell cycle entry. **(G)** Cell size at 20% budding for wild type and *TEF-WHI5* cells. Graph shows the average of three biological replicates. Errors bars are S.E.M.

To test the effects of a larger increase in Whi5 protein levels, we put *WHI5-3xHA* under the control of the *TEF1* promoter. In rich carbon, *TEF1-WHI5-3xHA* produced 12-fold more protein than the endogenous promoter, but caused only a modest effect on cell size (**Figures 3B,C**). In poor carbon, the *TEF1* promoter produced 10-fold more Whi5-3XHA and caused a slightly larger increase in cell size compared to rich carbon. The stronger effects of *TEF1-WHI5-3xHA* on cell size in poor carbon could again be due to the large decrease in Cln3 levels in poor carbon. *TEF1-WHI5-3xHA* caused greater effects on the size of *bck2Δ* cells (**Figure 3 – figure supplement 1C**). Finally, we found that expression of *WHI5-3XHA* from the *GAL1* promoter caused a 23-fold increase in Whi5 protein levels but only a modest increase in cell size (**Figure 3 – figure supplement 1D**). Several previous studies observed stronger effects of *GAL1-WHI5* on cell size (de Bruin *et al.* 2004; Barber *et al.* 2020).

We next directly tested the effects of *TEF1-WHI5-3XHA* on cell size at cell cycle entry in G1 phase. To do this, we grew *TEF1-WHI5-3XHA CLN3-6XHA* cells in poor carbon and used centrifugal elutriation to isolate small unbudded cells in early G1 phase. The cells were then released into rich or poor carbon and levels of Cln3-6XHA and Whi5-3XHA were analyzed by western blot (**Figure 3D**). The Whi5 dilution model requires that no new Whi5 be produced during growth in G1 phase (Schmoller *et al.* 2015). Here, we found that the *TEF1* promoter drove a gradual increase in Whi5 protein levels during G1, which would prevent dilution of Whi5 (**Figures 3D and Figure 3 – supplement 1E**). However, *TEF1-WHI5-3XHA* had little effect on cell size at cell cycle entry in rich or poor carbon (**Figures 3E,F,G)**. A recent study that utilized a different heterologous promoter to control Whi5 production also concluded that synthesis of Whi5 during G1 phase has little effect on cell size (Barber *et al.* 2020).

Since Whi5 is diluted by less than 50% during growth, one would expect a 2-fold increase in Whi5 to have a substantial effect on cell size if Whi5 dilution plays an important role in cell size control. The fact that changes in Whi5 protein levels of 10-fold or greater have modest effects on cell size, combined with the finding that synthesis of Whi5 during growth also causes modest effects, suggests that growth-dependent changes in Whi5 concentration are unlikely to play a major role in cell size control.

### Blocking membrane trafficking events required for cell growth prevents accumulation of Cln3

Together, the preceding experiments suggest that changes in the concentration of Whi5 do not play a substantial role in mechanisms that link cell cycle entry to cell growth. Rather, the data are more consistent with a model in which accumulation of Cln3 could be the critical readout of growth that triggers cell cycle entry. We therefore next investigated the relationship between accumulation of Cln3 and cell growth. As a first step, we sought to test whether accumulation of Cln3 protein in G1 phase is dependent upon growth. To do this, we first searched for ways to enforce a rapid block to growth during G1 phase. In budding yeast, vesicular traffic that drives bud growth occurs along actin cables and depolymerization of actin causes rapid cessation of bud growth. However, we discovered that addition of the actin depolymerizing drug latrunculin A and the microtubule depolymerizing drug nocodazole had no effect on growth during G1 phase, even though bud emergence was completely blocked (**Figures 4 A,B**).

**Figure 4:**
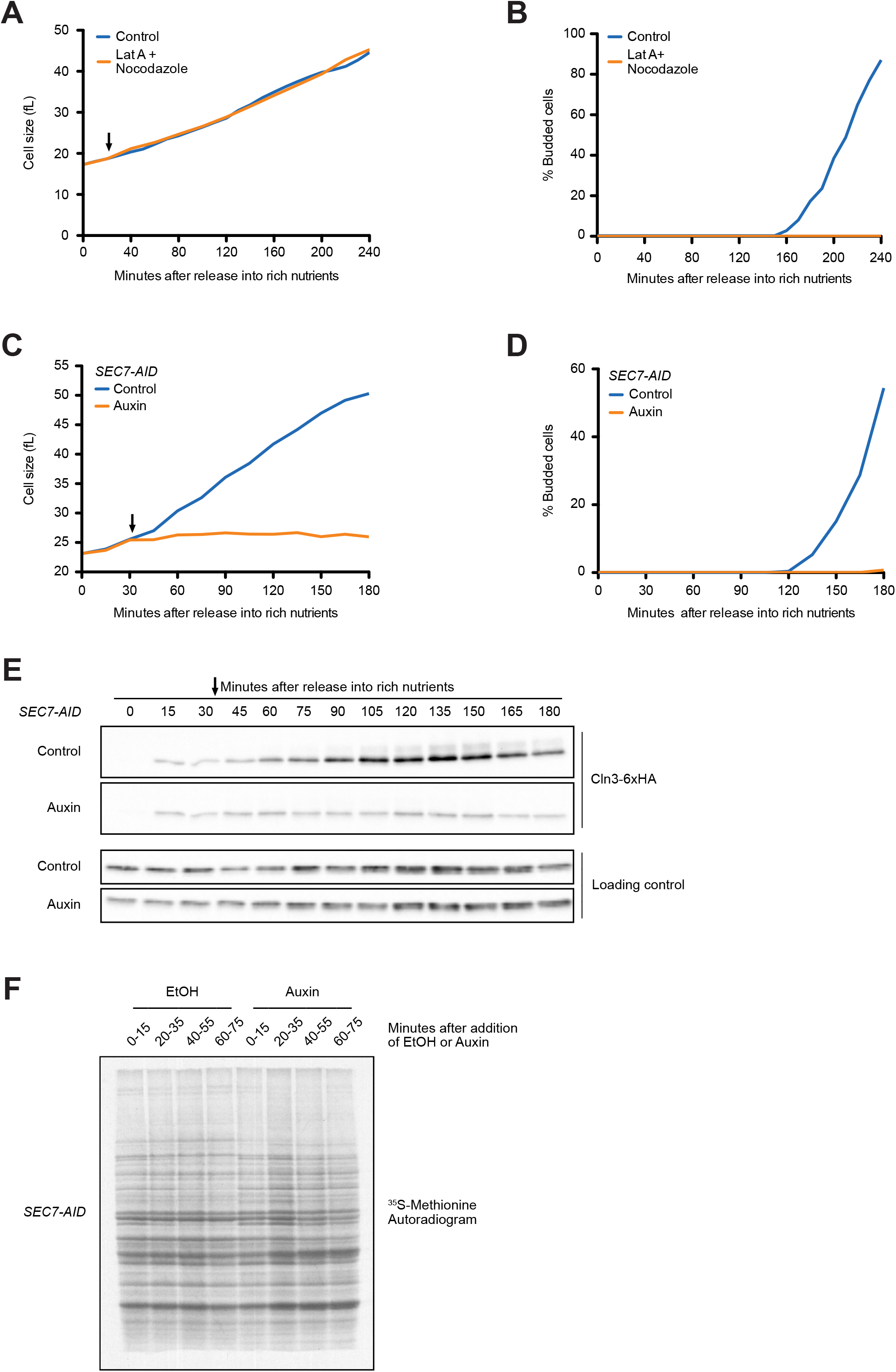
**(A-B)** Wild type cells were grown to mid-log phase in poor carbon (YPG/E) and small unbudded cells were isolated by centrifugal elutriation. The cells were split into two cultures and were then released into rich carbon (YPD) at 25°C. After 20 min, 100 μM of Latrunculin A and 20 μM nocodazole were added to one culture (arrow). **(A)** Median cell size was measured using a Coulter Counter and plotted as a function of time. **(B)** The percentage of budded cells as a function of time. **(C-E)***SEC7-AID* cells were grown to mid log phase in poor carbon and small unbudded cells were isolated by centrifugal elutriation. The cells were split into two cultures and were then released into rich carbon at 25°C. After 30 min, 1 mM auxin was added to one culture (arrow). **(C)** Median cell size was measured using a Coulter Counter and plotted as a function of time. **(D)** The percentage of budded cells as a function of time. **(E)** The behavior of Cln3-6xHA in the *sec7-AID* cells was analyzed by western blot. An anti-Nap1 antibody was used for a loading control. **(F)** Autoradiogram of ^35^S-methionine labeling to detect *de novo* protein synthesis. *SEC7-AID* cells were grown to mid log phase in-MET synthetic media containing glucose. After auxin or vehicle addition, 1.2 ml samples collected from the cultures were labeled with ^35^S-methionine for 15 min intervals starting every 20 min to measure the rate of protein synthesis within the 15 min intervals.

We next tested whether proteins that drive early steps in membrane trafficking pathways are required for growth in G1 phase. Sec7 is required for ER-to-Golgi transport (Lupashin *et al.* 1996; Deitz *et al.* 2000). Inactivation of Sec7 using an auxin-inducible degron (sec7-AID) caused a rapid and complete block of growth in G1 phase (**Figure 4C**), as well as a complete block of bud emergence (**Figure 4D**). Inactivation of Sec7 completely blocked the gradual accumulation of Cln3 protein that normally occurs during growth in G1 phase (**Figure 4E**). We obtained similar results using a temperature sensitive allele of SEC7 (*sec7-1*) (not shown).

One explanation for the effects of inactivating Sec7 could be that blocking membrane traffic causes a general decrease in the rate of protein synthesis, which leads to a failure to accumulate Cln3. However, previous work found that protein synthesis continues after membrane traffic is blocked (Novick and Schekman 1979). Moreover, we found that levels of a loading control protein increased normally, which argues against a general decrease in the rate of protein synthesis (**Figures 4E and Figure 4 – figure supplement 1**). Finally, we directly measured the rate of protein synthesis in *SEC7-AID* cells via incorporation of ^35^S-methionine. To do this, auxin was added to asynchronously growing *SEC7-AID* cells and the rate of incorporation of ^35^S-methionine was measured during 15 minute intervals before and after addition of auxin (**Figure 4F**). We detected a few changes in the pattern of proteins synthesized in the *SEC7-AID* cells, which is likely caused by a failure in post-translational processing of proteins in the secretory pathway. However, we did not detect a decrease in the rate of protein synthesis. We obtained similar results when we induced destruction of *SEC7-AID* in synchronized cells undergoing growth in G1 phase (not shown). These data suggest that failure to accumulate Cln3 when Sec7 is inactivated is not due to a general decrease in the rate of protein synthesis.

Together, these observations show that the gradual rise in Cln3 levels during G1 phase is dependent upon membrane trafficking events that drive plasma membrane growth, consistent with the idea that Cln3 levels could provide a readout of cell growth.

### The budding yeast homologs of mammalian SGK kinases are required for accumulation of Cln3 during G1 phase

To further explore the relationship between cell growth and gradual accumulation of Cln3, we searched for signals that influence cell growth and size in G1 phase, with the goal of testing whether these signals also influence Cln3 accumulation. Polar bud growth that occurs after G1 phase requires cyclin-dependent kinase (CDK) activity (McCusker *et al.* 2007). We therefore first tested whether CDK activity is required for growth in G1 phase. There are two cyclin-dependent kinases that play overlapping roles in G1 phase: Cdk1 and Pho85. We utilized a strain that is dependent upon analog-sensitive alleles of both kinases (*cdk1-as pho85-as*) to test whether cyclin-dependent kinases are required for growth in G1 phase. Inhibition of both kinases had no effect on growth in G1 phase (**Figure 5 – figure supplement 1**). This observation, along with discovery that actin is not required for growth in G1 phase (**Figure 4A**), suggests that there may be substantial differences between the mechanisms that drive growth in G1 phase and those that drive growth of the daughter bud.

We next tested Tor kinase signaling pathways, which play conserved roles in control of cell growth. Tor kinases are assembled into two distinct multi-protein signaling complexes called TORC1 and TORC2 (Loewith *et al.* 2002). A key downstream target of TORC1 is the Sch9 kinase, a member of the AGC kinase family that is thought to be the functional ortholog of vertebrate S6 kinase. Sch9 mediates TORC1-dependent control of ribosome biogenesis (Urban *et al.* 2007). Cells that lack Sch9 are viable but proliferate slowly and show a large decrease in cell size (Jorgensen *et al.* 2002; Jorgensen and Tyers 2004a). Inhibition of an analog-sensitive allele of *SCH9* (*sch9-*as) had no effect on growth rate (**Figure 5A**) but caused a slight delay in bud emergence (**Figure 5B**). Inhibition of sch9-as also caused a reduction in Cln3 levels but did not block gradual accumulation of Cln3 during growth in G1 phase (**Figure 5C**).

**Figure 5:**
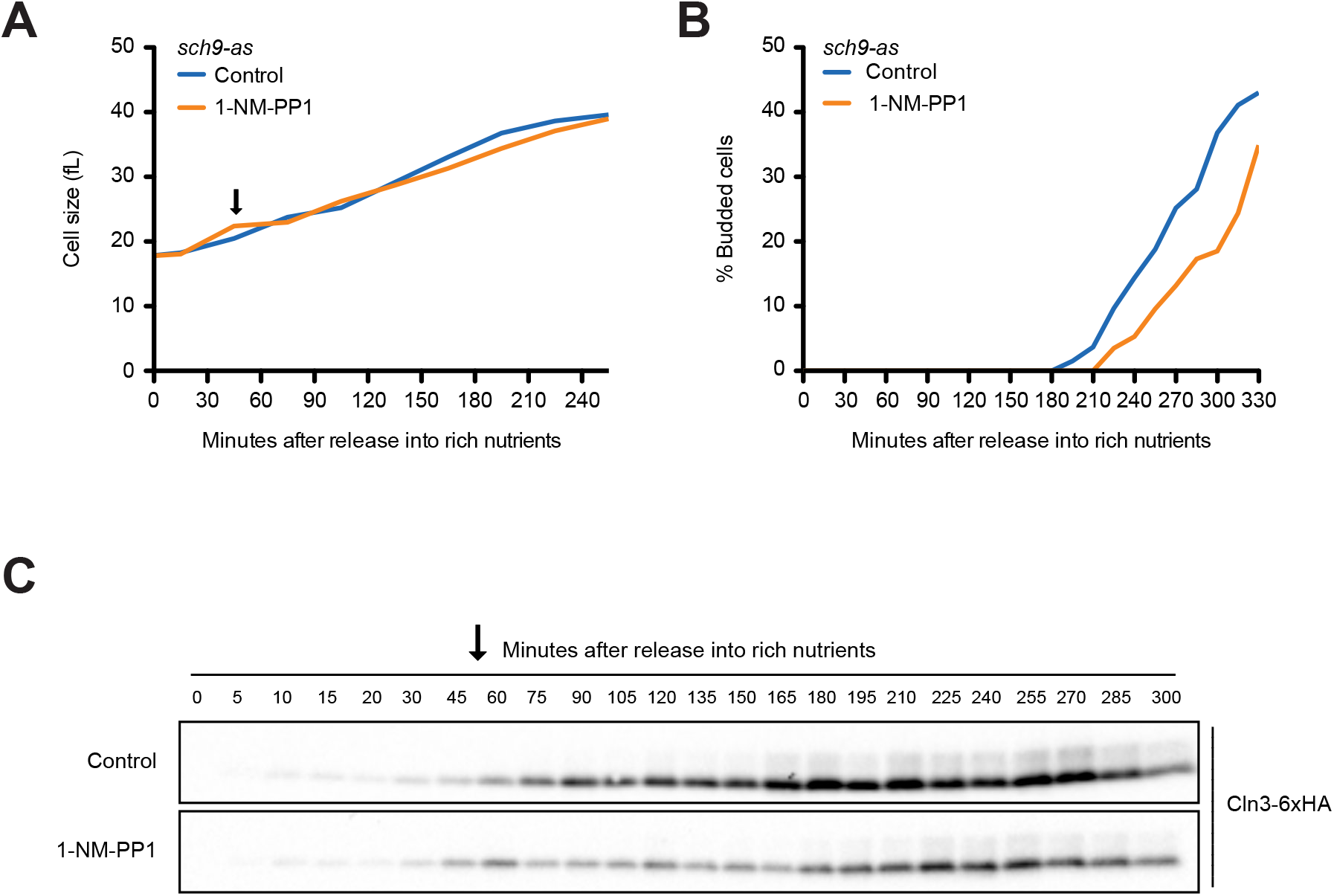
*sch9-as* cells were grown to mid log phase in poor carbon (YPG/E) and small unbudded cells were isolated via centrifugal elutriation. The cells were split into two cultures and were released into rich carbon (YPD, not supplemented with additional adenine (see Methods)). After 45 min, 250 nM 1-NM-PP1 was added to one culture (arrow). **(A)** Median cell size was measured using a Coulter Counter and plotted as a function of time. **(B)** The percentage of budded cells as a function of time. **(C)** The behavior of Cln3-6xHA in the *sch9-as* cells was analyzed by western blot. An anti-Nap1 antibody was used for a loading control.

We next tested whether components of a TORC2 signaling network are required for growth in G1 phase. A key function of TORC2 is to activate a pair of redundant kinase paralogs called Ypk1 and Ypk2, which are the budding yeast homologs of vertebrate SGK kinases (Casamayor *et al.* 1999; Kamada *et al.* 2005). Previous work has shown that Ypk1/2 are required for normal control of cell growth and size (Lucena *et al.* 2018). For example, loss of Ypk1 causes a large decrease in cell size, as well as a reduced rate of proliferation. Loss of both Ypk1 and Ypk2 is lethal.

To test whether Ypk1/2 influence cell growth and accumulation of Cln3 in G1 phase, we utilized an analog-sensitive allele of *YPK1* in a *ypk2Δ* background (*ypk1-as ypk2Δ*) (Sun *et al.* 2012). Inhibition of ypk1-as in early G1 phase caused a decrease in growth rate and a complete failure in bud emergence (**Figures 6A,B**). It also caused a rapid and complete loss of Cln3 protein (**Figure 6C**). In wild type control cells, the inhibitor caused a slight decrease in growth rate but had little effect on Cln3 levels (**Figure 6 – figure supplement 1A**). The loss of Cln3 after inhibition of ypk1-as did not appear to be caused by a general shutdown of protein synthesis because growth continued and levels of a loading control protein increased (**Figures 6A, 6C and Figure 6 – figure supplement 1C**). Direct measurement of the rate of protein synthesis confirmed that inhibition of ypk1-as did not cause a general shutdown of protein synthesis (**Figure 6 – figure supplement 1D**). Previous work has shown that *ypk1Δ* alone causes a large decrease in cell size (Lucena et al., 2018). Here, we found *ypk1Δ* caused a large reduction in Cln3 levels in asynchronous cells in both rich and poor carbon (**Figure 6D**).

**Figure 6:**
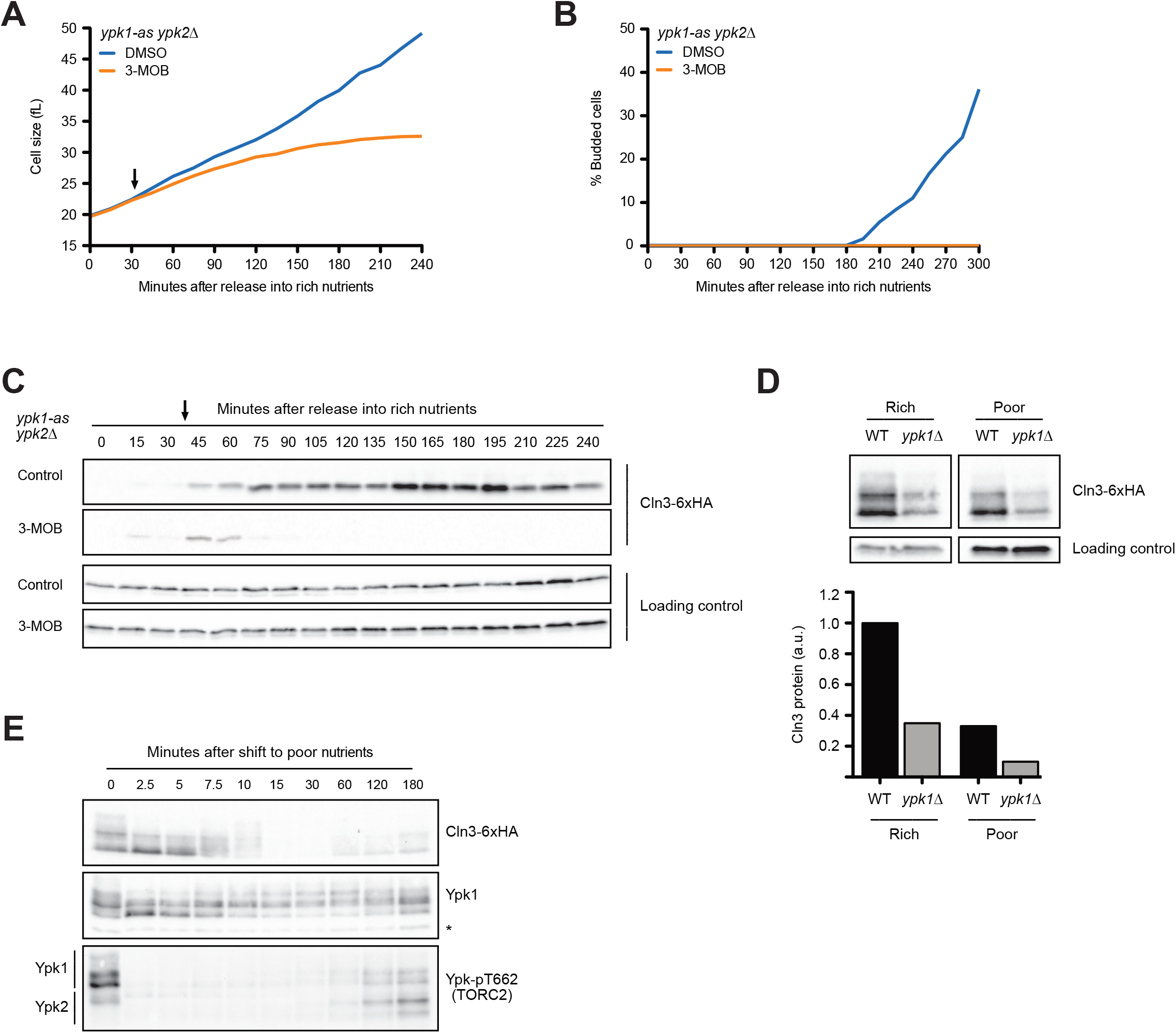
**(A-C)** *ypk1-as ypk2Δ* cells were grown overnight in poor carbon (YPG/E) to mid log phase and small unbudded cells were isolated by centrifugal elutriation. The cells were divided into two cultures and were then released into rich carbon (YPD, not supplemented with additional adenine (see Methods)). After 30 min 25 μM 3-MOB-PP1 was added to one culture. **(A)** Median cell size was measured using a Coulter counter and plotted as a function of time. **(B)** The percentage of budded cells as a function of time. **(C)** The behavior of Cln3-6xHA in the *ypk1-as ypk2Δ* cells was analyzed by western blot. An anti-Nap1 antibody was used for a loading control. Arrow indicates when 3-MOB-PP1 was added. **(D)** Wild type and *ypk1Δ* cells were grown to mid log phase in rich or poor carbon. The levels of Cln3-6xHA protein were analyzed by western blot. An anti-Nap1 was used for a loading control. Levels of Cln3-6XHA protein were quantified relative to levels of Cln3-6XHA in wild type cells in rich carbon. Cln3 protein levels were first normalized to the loading control. **(E)** Wild type cells were grown to mid log phase in rich carbon and then shifted to poor carbon at 25°C. The behavior of Cln3-6xHA, TORC2-dependent phosphorylation of Ypk1/2 and Ypk1 protein were assayed by western blot. A phosphospecific antibody was used to detect TORC2-dependent phosphorylation of Ypk1/2 on T662 (ref). The antibody detects phosphorylated forms of both Ypk1 and Ypk2. Total Ypk1 protein was detected with an anti-Ypk1 antibody. Asterisk indicates a background band that also serves as a loading control.

Ypk1/2 undergo complex regulation and are phosphorylated by multiple kinases. We found that changes in Cln3 protein levels that occur during a shift from rich to poor carbon closely paralleled changes in the phosphorylation state of Ypk1 that can be detected via shifts in electrophoretic mobility (**Figure 6E**). Thus, the transient increase in Cln3 levels immediately after a shift to poor carbon was correlated with a loss of Ypk1 phosphorylation, and the decrease in Cln3 levels starting at 10 minutes was correlated with an increase in Ypk1 phosphorylation. A reappearance of Cln3 at the end of the time course was accompanied by another decrease in Ypk1 phosphorylation. The phosphorylation of Ypk1 that can be detected by electrophoretic mobility shifts appears to be due at least partly to related kinase paralogs called Fpk1 and Fpk2; however, the functions of these phosphorylation events are poorly understood (Roelants *et al.* 2010). Nevertheless, the close correlation between changes in Ypk1 phosphorylation and changes in Cln3 protein levels provide further evidence for a connection between Ypk1 signaling and Cln3 protein levels.

TORC2-dependent phosphorylation of Ypk1 and Ypk2 can be detected with a phospho-specific antibody that recognizes a site found on both kinases (referred to as T662 in Ypk1) (Niles *et al.* 2012). Previous studies have shown that a shift from rich to poor carbon causes rapid loss of TORC2-dependent phosphorylation of Ypk1/2 (Lucena *et al.* 2018). Here, we found that loss of TORC2-dependent phosphorylation of Ypk1/2 upon a shift to poor carbon was immediate and preceded the decrease in Cln3 levels by approximately 10 minutes **(Figure 6E, bottom panel)**. The increase in Cln3 that occurred later in the time course as cells adapted to the new carbon source was correlated with an increase in TORC2-dependent phosphorylation of Ypk1/2. Thus, changes in Cln3 levels were partially correlated with TORC2-dependent phosphorylation of Ypk1/2.

Together, the data show that Ypk1/2, which were previously shown to play roles in control of cell growth and size, also play a role in regulating levels of Cln3. The data further suggest that Cln3 protein levels are unlikely to reflect a simple readout of translation rate. Rather, the data indicate that poorly understood signals associated with plasma membrane growth and the TORC2 network strongly influence Cln3 levels.

### Sphingolipid-dependent signals influence Cln3 levels

A key function of Ypk1/2 is to control the activity of a biosynthetic pathway that builds sphingolipids and ceramide lipids (Aronova *et al.* 2008; Roelants *et al.* 2011; Sun *et al.* 2012; Muir *et al.* 2014). A simplified overview of the role of Ypk1/2 in controlling synthesis of sphingolipids and ceramides is shown in **Figure 7 – figure supplement 1A**). Ypk1/2 stimulate the enzyme that initiates production of sphingolipids, and also stimulate the activity of ceramide synthase, which builds ceramide from sphingolipid precursors. Sphingolipids and ceramides play poorly understood roles in signaling. In previous work, we showed that Ypk1/2-dependent control of the ceramide synthesis pathway is required for normal control of cell growth and size (Lucena *et al.* 2018). For example, myriocin, a small molecule inhibitor of sphingolipid synthesis, causes a dose-dependent decrease in growth rate in G1 phase and a corresponding decrease in cell size at cell cycle entry. Similarly, loss of ceramide synthase causes a large reduction in cell size, as well as a complete failure in nutrient modulation of cell size, growth rate, and TORC2 activity.

To test whether Ypk1/2 control Cln3 levels via sphingolipids or ceramides, we first tested the effects of myriocin, which inhibits the first step in production of sphingolipids. In previous work, we analyzed the effects of sub-lethal doses of myriocin on cell growth and size in G1 phase (Lucena *et al.* 2018). Here, we used a higher concentration of myriocin that blocks proliferation. Addition of myriocin caused a large reduction in growth rate in G1 phase and delayed bud emergence but caused only a modest decrease in Cln3 levels (**Figures 7A-C**). Myriocin also caused cells to initiate bud emergence at a smaller size (**Figure 7D**). Thus, the complete loss of Cln3 caused by inhibition of Ypk1/2 is not due to a loss of sphingolipid-dependent signals.

**Figure 7:**
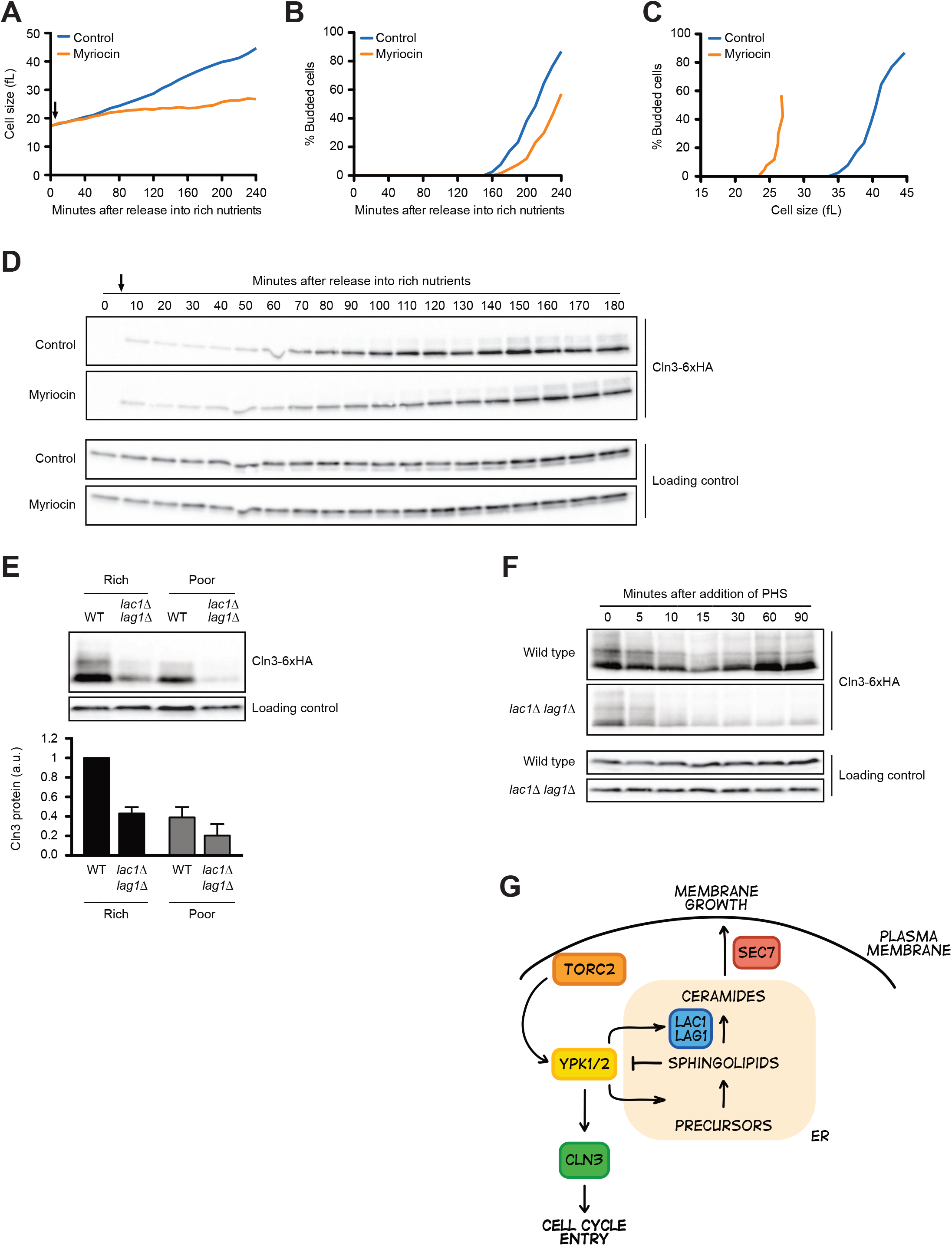
**(A-D)** Wild type cells were grown overnight in poor carbon (YPG/E) and small unbudded cells were isolated via centrifugal elutriation. The cells were split into two cultures and released into rich carbon (YPD) at 25°C. After taking the initial time point, 5 μg/ml myriocin was added to one culture (arrow). **(A)** Median cell size was measured using a Coulter counter and plotted as a function of time. **(B)** The percentage of budded cells as a function of time. **(C)** The percentage of budded cells plotted as a function of cell size at each time point. **(D)** The levels of Cln3-6xHA protein were analyzed by western blot. An anti-Nap1 antibody was used for a loading control. **(E)** Wild type and *lac1Δ lag1Δ* cells were grown to mid log phase in rich and poor carbon. The levels of Cln3-6xHA protein were analyzed by western blot. An anti-Nap1 antibody was used for a loading control. Levels of Cln3-6XHA protein were quantified relative to levels of Cln3-6XHA in wild type cells in rich carbon. Cln3 protein levels were first normalized to the loading control. **(F)** Wild type and *lac1Δ lag1Δ* cells were grown to mid log phase in rich carbon (YPD). 20 μM phytosphingosine (PHS) was added to each culture and the cultures were incubated at 25°C. The levels of Cln3-6xHA protein were analyzed by western blot. An anti-Nap1 antibody was used for a loading control. **(G)** A model depicting how Cln3 levels could be modulated by Ypk1/2.

In previous work, we found that ceramide-dependent signals modulate TORC2 activity via a feedback loop and are also required for normal control of cell growth and size (Lucena *et al.* 2018; Alcaide-Gavilan *et al.* 2018). We therefore also tested the effects of inactivating ceramide synthase, which builds ceramide from sphingolipid precursors. The catalytic subunit of ceramide synthase is encoded by a pair of redundant paralogs called *LAC1* and *LAG1* (**Figure 7 – figure supplement 1A)**. In contrast to myriocin, we found that *lac1Δ lag1Δ* caused a substantial decrease in levels of Cln3 in asynchronous cells (**Figure 7E**). This was surprising, because both myriocin and inactivation of ceramide synthase block production of ceramide, yet they appear to have different effects on Cln3. An explanation could be that inactivation of ceramide synthase leads to a build-up of sphingolipids that inhibit production of Cln3. To test this, we added exogenous sphingolipids to cells. In previous work, we found that addition of the sphingolipid phytosphingosine to cells causes a rapid and dramatic decrease in TORC2 signaling to Ypk1/2 (Lucena *et al.* 2018). The decrease in TORC2 signaling was dependent upon ceramide synthase, which provided evidence for ceramide-dependent feedback signaling to TORC2. Here, we found that exogenously added phytosphingosine caused a rapid loss of Cln3 in wild type cells (**Figure 7F**). Levels of Cln3 recovered in wild type cells, most likely due to conversion of the added phytosphingosine to ceramide. In contrast, Cln3 levels did not recover in *lac1Δ lag1Δ* cells.

These data show that sphingolipid-dependent signals strongly influence Cln3 levels, and provide further evidence that complex signaling mechanisms control accumulation of Cln3 protein. Models that could explain these observations are considered in the Discussion.

## Discussion

### The concentration of Cln3 protein increases during growth in G1 phase

One class of models for cell size control suggests that accumulation of Cln3 is a key reporter of cell growth during G1 phase (Jorgensen and Tyers 2004b; Turner *et al.* 2012). However, there have been conflicting reports about the behavior of Cln3 in G1 phase (Zapata *et al.* 2014; Schmoller *et al.* 2015; Litsios *et al.* 2019). Here, we used quantitative western blotting to analyze the behavior of Cln3 during growth in G1 phase in rich versus poor carbon. Under the growth conditions used for our experiments, we found that the number of Cln3 molecules per cell increases approximately 10-fold before cell cycle entry in rich carbon and 2.5 fold in poor carbon. When combined with growth data, these data suggest that Cln3 concentration increases 7-fold before bud emergence in rich carbon and 2-fold in poor carbon. The large increase in Cln3 concentration observed for cells growing in rich carbon may be due partially to the fact that the cells were grown in poor carbon prior to synchronization and therefore underwent an unusually extensive growth interval to reach the increased size characteristic of cells in rich carbon.

In both rich and poor carbon, synthesis of Cln3 protein outpaces the overall rate of protein synthesis that would be needed for the less than 2-fold increase in cell size that occurs during growth in the same interval. Thus, changes in Cln3 concentration could generate a signal with substantial dynamic range that could be used to trigger cell cycle entry. Similar conclusions were reached in a previous study that used an in vivo reporter to analyze Cln3 translation rates and targeted mass spectrometry to analyze Cln3 protein levels (Litsios *et al.* 2019).

### The threshold amount of Cln3 required for cell cycle entry is dramatically reduced in poor carbon

Simple models based on previous studies would suggest that cell cycle entry at a reduced cell size in poor carbon could be achieved by reducing the levels of Whi5 protein, or by increasing levels of Cln3. However, we found that Cln3 levels are reduced nearly 10-fold in poor carbon, while levels of Whi5 do not change. Thus, cells in poor carbon undergo cell cycle entry despite a nearly 10-fold decrease in the ratio of Cln3 to Whi5. We also found that Cln3 protein levels are exquisitely sensitive to changes in carbon source that influence growth rate. Thus, a shift from rich to poor carbon, which causes a rapid decrease in growth rate, causes a transient increase in Cln3 protein levels within 5 minutes, followed by a dramatic reduction in Cln3 levels. These data show that Cln3 is highly responsive to nutrient-dependent signals that control cell growth and size, as expected for a critical regulator of cell size in G1 phase.

One estimate suggests that there could be as few as 100 molecules of Cln3 per cell in rich carbon (Cross *et al.* 2002), which would suggest that there could be as few as 10-20 molecules of Cln3 per cell in poor carbon. In contrast, Whi5 could be present at approximately 2500 molecules per cell, although there is considerable variance in estimates of the abundance of Whi5 protein (Ho *et al.* 2018; Dorsey *et al.* 2018). The mechanism that sets the threshold amount of Cln3 required to overcome Whi5 inhibition to initiate cell cycle entry is key to understanding how cell cycle progression is linked to cell growth yet remains one of the central enigmas of cell size control.

### The concentration of Whi5 protein has little influence on cell size

To investigate the role of changes in Whi5 concentration in controlling cell cycle entry, we tested whether overexpression of *WHI5* influences cell size. We found that an extra copy of untagged Whi5 expressed from its own promoter had little effect on cell size in wild type cells growing in YP media containing rich or poor carbon. We also found that 12-fold overexpression of *WHI5-3XHA* from the *TEF1* promoter had only a modest effect on cell size in rich carbon. Overexpression of *WHI5-3XHA* in poor carbon caused a more substantial increase in cell size, which could be due to the large decrease in Cln3 levels in poor carbon. Overexpression of *WHI5-3XHA* by 23-fold from the *GAL1* promoter caused a larger increase in cell size but did not cause a lethal cell cycle arrest as might be expected for a critical dose-dependent inhibitor of G1 phase progression.

The effects of Whi5 overexpression on cell size has been the primary mechanistic test of the Whi5 dilution model; however, increased dosage of Whi5 could also influence a positive feedback loop in which the late G1 cyclins Cln1 and Cln2 promote cell cycle entry via inactivation of Whi5 (Cross and Tinkelenberg 1991). Thus, current data do not distinguish whether the effects of Whi5 overexpression could be due to inhibition of the initial Cln3-dependent inactivation of Whi5 versus inhibition of Cln1/2-dependent inactivation of Whi5 via feedback. A more stringent test of the model is to examine the effects of synthesizing Whi5 during G1 phase to prevent dilution. A previous study tested this by using a heterologous promoter to drive Whi5 accumulation during G1 phase but did not observe the effects predicted by the Whi5 dilution model (Barber *et al.* 2020). Here, we found that Whi5 overexpressed 12-fold from the *TEF1* promoter was synthesized and accumulated gradually during G1 phase but did not cause major defects in cell size. We conclude that growth-dependent reduction in Whi5 is unlikely to play a major role in measuring cell size in G1 phase. Three recent studies reached similar conclusions (Dorsey *et al.* 2018; Litsios *et al.* 2019; Barber *et al.* 2020).

Dilution models for cell size control present a number of unresolved mechanistic issues. For example, dilution models require that key factors show a concentration-dependent activity. The only known activity of Whi5 is to bind and inhibit the SBF transcription factor. For Whi5 to repress transcription at SBF promoters, it must be present in excess over SBF binding sites and bind with sufficient affinity to achieve high occupancy at promoters. Moreover, to achieve high occupancy at SBF promoters Whi5 must be present at a concentration higher than the K_d_ for its interaction with SBF. One study estimated the concentration of Whi5 in the nucleus to be 120 nM (Dorsey *et al.* 2018). Independent studies that measured the total number of Whi5 molecules in the cell and nuclear volume suggest that the concentration could be higher (Jorgensen *et al.* 2007; Ho *et al.* 2018). The Kd for the interaction between Whi5 and SBF is unknown but could easily be in the low nM range. For comparison, the Rb transcriptional inhibitor binds to the E2F transcription factor with a K_d_ of 40 nM (Burke *et al.* 2010). Thus, the concentration of Whi5 in the nucleus may be substantially higher than the K_d_ for binding to SBF. In this case, a less than 2-fold change in cell volume would have little effect on occupancy of Whi5 at SBF promoters, which is the only known activity of Whi5. It is proposed that growth-dependent dilution causes a change in the ratio of Cln3 to Whi5. However, under the poor carbon conditions used to test the dilution model, the concentration of Whi5 is clearly much higher than the concentration of Cln3, so a less than 2-fold change in Whi5 concentration will have little effect on the ratio. Phosphorylation of Whi5 by Cdk1 could reduce the affinity of Whi5 for SBF, but in that case there may be no reason to hypothesize that anything other than rising Cln3 levels lead to inactivation of Whi5. It is certainly possible that there are biochemical parameters for the mechanisms that inactivate Whi5 that would be compatible with dilution models. However, in the absence of any data on those parameters it remains unclear whether dilution models are viable.

### Accumulation of Cln3 during G1 phase is dependent upon membrane trafficking events that drive cell growth

Previous studies have proposed that accumulation of Cln3 protein could be a key readout of cell growth. To provide a measure of growth, accumulation of Cln3 must be dependent upon growth. To test this, we used an auxin-inducible degron allele of *SEC7* to block membrane trafficking events that drive cell growth. Inactivation of *SEC7* in early G1 phase blocked cell growth, as well as accumulation of Cln3. This observation, combined with the results of our analysis of Cln3 accumulation in wild type cells, suggests that gradual accumulation of Cln3 in G1 phase is dependent upon growth and correlated with the extent of growth, as expected for a protein that provides a readout of growth.

In previous work we found additional evidence that membrane trafficking events that drive cell growth generate signals that are correlated with growth, which could be used to measure growth. For example, delivery of vesicles to the plasma membrane in growing daughter buds generates signals that lead to phosphorylation of the kinase Pkc1 and two related kinases called Gin4 and Hsl1 (Anastasia *et al.* 2012; Jasani *et al.* 2020). In each case, multi-site phosphorylation of these kinases occurs gradually during bud growth, and phosphorylation appears to be dependent upon growth and the extent of phosphorylation is correlated with the extent of growth. The signals relayed by these kinases are required for normal regulation of the duration and extent of bud growth; however, there is no evidence that they influence growth in G1 phase.

### Cln3 protein levels are controlled by the budding yeast homologs of mammalian SGK kinases

We also searched for signals that drive growth-dependent accumulation of Cln3. Growth of the daughter bud after cell cycle entry is dependent upon Cdk1 activity (McCusker *et al.* 2007). However, growth in G1 phase and accumulation of Cln3 showed no dependence on Cdk1 activity. Surprisingly, growth in G1 phase was also not dependent upon actin filaments, which are required for bud growth. The fact that bud growth is strongly dependent upon Cdk1 and actin filaments, whereas growth in G1 phase is not, suggests that there may be substantial differences in the mechanisms that drive cell growth at different stages of the cell cycle.

Inhibition of the protein kinase Sch9, a key downstream target of TORC1 that controls ribosome biogenesis, had little effect on the rate or extent of growth in G1 phase and caused relatively minor defects in accumulation of Cln3. This was surprising because *sch9Δ* causes a large decrease in both growth rate and cell size (Jorgensen and Tyers 2004a). Our results suggest that these defects could be a long term consequence of a decreased rate of ribosome biogenesis. For example, it is possible that cells in early G1 phase have sufficient ribosome capacity to get through the early stages of the cell cycle so that strong effects of loss of Sch9 only become apparent when the existing pool of ribosomes becomes insufficient as more growth occurs.

Inhibition of Ypk1/2, the budding yeast homologs of human SGK kinases, caused a rapid and complete loss of Cln3 protein. Previous studies have shown that Ypk1/2 relay nutrient-dependent signals and are required for normal control of cell growth and size (Lucena *et al.* 2018). Thus, the discovery that Ypk1/2 control Cln3 levels establishes a link between Ypk1/2 and Cln3 that could help explain how Ypk1/2 and nutrient-dependent signals modulate cell size. Further investigation of the mechanisms that link Cln3 levels to Ypk1/2 activity should provide important clues to how cell growth and size are controlled.

Ypk1/2 are direct targets of TORC2 and TORC2 signaling is strongly modulated by carbon source, which suggests that TORC2 could play a role in modulating Cln3 levels (Kamada *et al.* 2005; Niles and Powers 2012; Lucena *et al.* 2018; Alcaide-Gavilan *et al.* 2018). However, there are additional kinases that phosphorylate Ypk1/2 and regulation of Ypk1/2 remains poorly understood. (Casamayor *et al.* 1999; Roelants *et al.* 2010). Thus, it is also possible that there are signals that act in parallel with TORC2 to modulate Cln3 levels via Ypk1/2. A further complication is that Ypk1/2 relay poorly understood feedback signals that regulate their upstream kinases, which can make it difficult to clearly delineate direct versus indirect effects of manipulating the signaling network (Roelants *et al.* 2010; 2011; Berchtold *et al.* 2012; Lucena *et al.* 2018; Alcaide-Gavilan *et al.* 2018).

There was an important difference between the effects of inhibiting Ypk1/2 and the effects of inhibiting Sec7. Inhibition of Ypk1/2 caused a reduced rate of growth and a rapid and complete loss of Cln3. In contrast, inhibition of Sec7 completely blocked cell growth as well as further accumulation of Cln3 but did not cause a loss of Cln3. These observations suggest that the effects of inhibiting Ypk1/2 are not a simple consequence of a slowdown in overall growth, and that Ypk1/2 could play a more direct role in controlling Cln3 levels. A key function of Ypk1/2 is to stimulate production of sphingolipids and ceramides, which relay signals that influence cell growth and size. Myriocin, an inhibitor of the first step of sphingolipid synthesis, did not block accumulation of Cln3. Similarly, loss of ceramide synthase reduced but did not eliminate Cln3. Thus, the disappearance of Cln3 upon inhibition of Ypk1/2 is not due to a failure in synthesis of sphingolipids and ceramides. Rather, another target of Ypk1/2 likely exists that influences Cln3 levels.

The different effects of Sec7, Ypk1/2 and myriocin on Cln3 levels are puzzling. Previous studies found that myriocin, inhibition of Ypk1/2, and inhibition of membrane traffic all cause increased TORC2-dependent phosphorylation of Ypk1/2, which is thought to lead to increased activity of Ypk1/2 (Niles *et al.* 2014; Clarke *et al.* 2017; Lucena *et al.* 2018). In each case, the increased TORC2-dependent phosphorylation of Ypk1/2 is thought to be due to poorly understood negative feedback loops in the TORC2 network. We confirmed that each of these experimental manipulations also cause increased TORC2-dependent phosphorylation of Ypk1/2 during growth in G1 phase (not shown). Although each experimental manipulation causes a decrease in growth rate, they each have different effects on accumulation of Cln3. We also found that addition of exogenous sphingolipids to cells causes a rapid decline in Cln3 levels that is not dependent upon conversion of sphingolipids to ceramides.

What kind of model could explain these observations? One possibility is that there is a pool of Ypk1/2 at the endoplasmic reticulum that promotes accumulation of Cln3, and sphingolipids produced at the endoplasmic reticulum inhibit the Ypk1/2 that promotes Cln3 accumulation (**Figure 7G**). In this model, loss of Ypk1/2 activity would cause rapid loss of Cln3 because Ypk1/2 directly promote production of Cln3. Inactivation of Sec7 would lead to accumulation of sphingolipids in the endoplasmic reticulum because they fail to be transported to the Golgi apparatus, which would inhibit the Ypk1/2-dependent increase in Cln3 levels during cell growth. Myriocin would lead to loss of inhibitory sphingolipids at the endoplasmic reticulum so that Ypk1/2 can drive an increase in Cln3, even though growth has stopped. Finally, addition of exogenous sphingolipids would lead to an increase in sphingolipid-dependent signals that inhibit Ypk1/2, leading to loss of Cln3. Several previous studies have suggested that Cln3 is localized to the endoplasmic reticulum and that regulation of Cln3 is closely associated with endoplasmic reticulum function (Vergés *et al.* 2007; Yahya *et al.* 2013).

Data from previous studies are consistent with a model in which there are multiple differentially regulated pools of TORC2 and Ypk1/2. For example, Ypk1/2 must carry out functions at the endoplasmic reticulum because the Orm1/2 proteins and ceramide synthase, two well-established direct targets of Ypk1/2, are localized there. Yet TORC2 and Ypkl1/2 are also found at the plasma membrane (Berchtold and Walther 2009; Niles *et al.* 2012) and transport of sphingolipids or ceramide to the plasma membrane appears to be essential for relaying negative feedback signals that inhibit TORC2-dependent phosphorylation of Ypk1/2, consistent with the existence of signaling events at the plasma membrane (Clarke *et al.* 2017). Thus, there may be distinct, differentially regulated pools of Ypk1/2 at the endoplasmic reticulum and the plasma membrane.

At this point, numerous alternative models are possible, and our ability to construct detailed models is constrained by our limited understanding of the function and regulation of Ypk1/2. Nevertheless, the fact that Cln3, a critical dose-dependent regulator of cell size, is strongly influenced by signaling from key components of the TORC2 network suggests that a full understanding of cell size control will require a much better understanding of growth control and the TORC2 signaling network.

## Acknowledgments

We thank Andrew Murray, Jan Skotheim, Kurt Schmoller, Matthew Swaffer, Matthias Heinemann, Andreas Milias-Argeitis and members of the lab for helpful discussions. We also thank Jack Stevenson and Kevan Shokat for providing 3-MOB-PP1. This work was funded by NIH grants GM053959 and GM131826.

## METHODS

### Yeast strains and media

All strains are in the W303 background (*leu2-3, 112 ura3-1 can1-100 ade2-1 his3-11,15 trp1-1 GAL+ ssd1-d2*). All strains are derived from the parent strain DK186 (mat a) with the exception of DK2961, which is derived from Y3507 (mat α) (Gift from James R. Broach, Penn State). Table 1 shows additional genetic features. One-step, PCR-based gene replacement was used for making deletions and adding epitope tags at the endogenous locus(Longtine *et al.* 1998; Janke *et al.* 2004). A plasmid carrying a copy of WHI5 was made by amplifying the *WHI5* gene with 329bp upstream of the start codon and 159bp downstream of the stop codon from genomic DNA with oligos *GCGGGATCCCCGTCTTCCTTGTGCTGTTTATG* and *GCGGAATTCGCGGATGTTGATCGGCGGAT*. The PCR fragment was cloned into the BamH1 and EcoR1 sites of YIplac211 to create plasmid pRAS1. To integrate an extra copy of WHI5 into the genome the plasmid was cut with StuI to target integration at the *URA3* locus. The empty vector YIplac211 was integrated to create isogenic control strains.

**Table 1:**
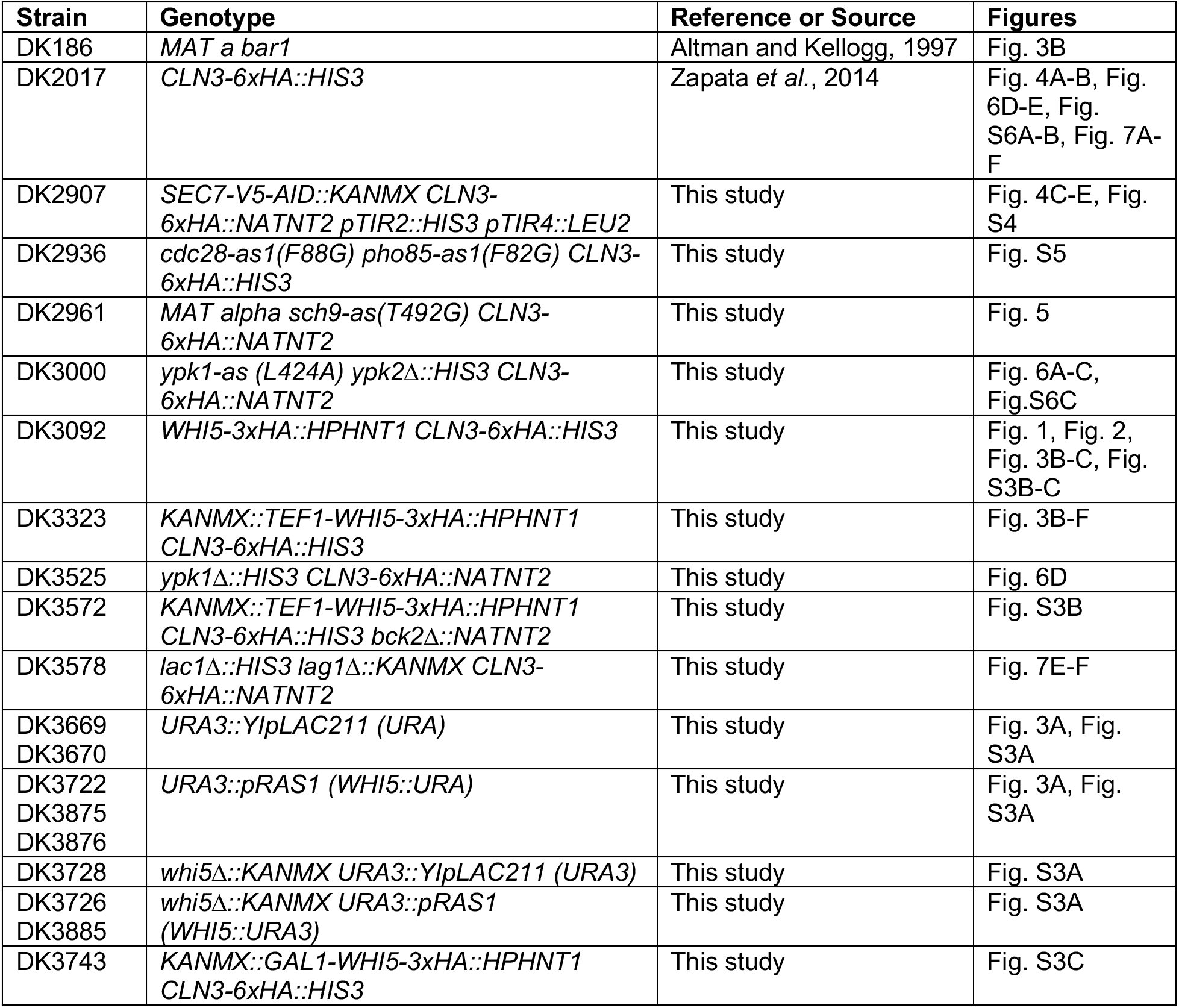

| **Strain** | **Genotype** | **Reference or Source** | **Figures** |
| --- | --- | --- | --- |
| DK186 | *MAT a bar1* | Altman and Kellogg, 1997 | Fig. 3B |
| DK2017 | *CLN3-6xHA::HIS3* | Zapata *et al.*, 2014 | Fig. 4A-B, Fig. 6D-E, Fig. S6A-B, Fig. 7A-F |
| DK2907 | *SEC7-V5-AID::KANMX CLN3-6xHA::NATNT2 pTIR2::HIS3 pTIR4::LEU2* | This study | Fig. 4C-E, Fig. S4 |
| DK2936 | *cdc28-as1(F88G) pho85-as1(F82G) CLN3-6xHA::HIS3* | This study | Fig. S5 |
| DK2961 | *MAT alpha sch9-as(T492G) CLN3-6xHA::NATNT2* | This study | Fig. 5 |
| DK3000 | *ypk1-as (L424A) ypk2Δ::HIS3 CLN3-6xHA::NATNT2* | This study | Fig. 6A-C, Fig. S6C |
| DK3092 | *WHI5-3xHA::HPHNT1 CLN3-6xHA::HIS3* | This study | Fig. 1, Fig. 2, Fig. 3B-C, Fig. S3B-C |
| DK3323 | *KANMX::TEF1-WHI5-3xHA::HPHNT1 CLN3-6xHA::HIS3* | This study | Fig. 3B-F |
| DK3525 | *ypk1Δ::HIS3 CLN3-6xHA::NATNT2* | This study | Fig. 6D |
| DK3572 | *KANMX::TEF1-WHI5-3xHA::HPHNT1 CLN3-6xHA::HIS3 bck2Δ::NATNT2* | This study | Fig. S3B |
| DK3578 | *lac1Δ::HIS3 lag1Δ::KANMX CLN3-6xHA::NATNT2* | This study | Fig. 7E-F |
| DK3669<br>DK3670 | *URA3::YlpLAC211 (URA)* | This study | Fig. 3A, Fig. S3A |
| DK3722<br>DK3875<br>DK3876 | *URA3::pRAS1 (WHI5::URA)* | This study | Fig. 3A, Fig. S3A |
| DK3728 | *whi5Δ::KANMX URA3::YlpLAC211 (URA3)* | This study | Fig. S3A |
| DK3726<br>DK3885 | *whi5Δ::KANMX URA3::pRAS1 (WHI5::URA3)* | This study | Fig. S3A |
| DK3743 | *KANMX::GAL1-WHI5-3xHA::HPHNT1 CLN3-6xHA::HIS3* | This study | Fig. S3C |

Cells were grown in YP medium (1% yeast extract, 2% peptone) that contained 40 mg/L adenine and a carbon source, except where noted in the figure legends. Rich carbon medium (YPD) contained 2% dextrose, while poor carbon medium (YPG/E) contained 2% glycerol and 2% ethanol. In experiments using the ATP analog inhibitors 1-NM-PP1, 1-NA-PP1, or 3-MOB-PP1, no additional adenine was added to the media. Complete synthetic media (CSM) (MP Biosciences) contained either 2% dextrose (rich carbon) or 2% glycerol and 2% ethanol (poor carbon) with an additional 40mg/L adenine. CSM without methionine was used for 35S-methionine labeling experiments.

Myriocin (Sigma) was solubilized in 100% methanol to make a 500 μg/mL stock solution. We have observed significant batch-to-batch differences in the effective concentration of myriocin from the same supplier. All ATP analog inhibitors were solubilized in 100% DMSO. 3-MOB-PP1 was a gift from the Shokat lab (UCSF). Auxin (indole-3-acetic acid) (Aldrich) stock was prepared at 50 mM in 100% EtOH and used at 1mM.

### Cell size analysis and bud emergence

Cells were grown in YPD or YPG/E medium overnight at 22°C to mid-log phase (OD_600_ less than 0.7). Cells were fixed with 3.7% formaldehyde for 30 min and were then washed with PBS + 0.02% Tween-20 + 0.1% sodium azide (PBTA) before measuring cell size using a Z2 Coulter Channelyzer as previously described (Lucena *et al.* 2018) using Z2 AccuComp v3.01a software. For log phase cultures, each cell size plot is an average of three independent biological replicates in which each biological replicate is the average of three technical replicates. For 2xWHI5 measurements, three independent strain isolates generated by integration of the WHI5 plasmid pRAS1 were measured in more than three independent biological replicates and averaged. Similarly, the isogenic control strain integrated with empty vector is an average of two independent isolates measured in more than three independent biological replicates and averaged. The percentage of budded cells was calculated by counting >200 cells at each time point using a Zeiss Axioskop 2 (Carl Zeiss).

### Isolation of small, unbudded cells by centrifugal elutriation

Strains were grown in YPG/E medium overnight to OD600 0.4-0.8 at 30°C except for strains harboring temperature-sensitive alleles and the *ypk1-as ypk2Δ* strain, which were grown overnight at 22°C. Cells were harvested by spinning at 4,000rpm in a JLA 8.1 rotor at 4°C for six minutes. Cell pellets were resuspended in ~100ml cold YPG/E, then sonicated for one minute at duty cycle 0.5 using a Braun-Sonic U sonicator with a Braun 2000U probe at 4°C. Cells were loaded onto a Beckman JE-5.0 elutriator in a Beckman Coulter J6-MI centrifuge spinning at 2,900rpm at 4°C. After 10 minutes of equilibration, small, unbudded cells were collected and then pelleted by spinning in a JA-14 rotor at 5,000rpm for five minutes. Cells were resuspended into fresh medium at OD600 0.4-0.6 and grown at 25°C in a shaking water bath. Samples were collected and fixed with 3.7% formaldehyde for 15-30 minutes for cell size measurements and to calculate the percentage of budded cells. Median cell size was calculated by the Coulter AccuComp software for each time point. We found that cell size measurements of *ypk1-as ypk2Δ* cells isolated from centrifugal elutriation were more consistent when cells were not fixed with formaldehyde but were measured live after washing once with PBTA to remove YP media.

### Western Blotting

For western blotting, cells 1.6 ml samples taken from cultures were pelleted in a microfuge at 13,200rpm for 15 sec before aspirating the supernatant and adding 250 μL of glass beads and freezing on liquid nitrogen. Cells were lysed in 140 μL of 1X SDS-PAGE sample buffer (65 mM Tris-HCl, pH 6.8, 3% SDS, 10% glycerol, 100 mM β-glycerophosphate, 50 mM NaF, 5% β-mercaptoethanol, 2 mM PMSF, and bromophenol blue) by bead beating in a Biospec Mini-Beadbeater-16 at 4°C for two minutes. The lysate was centrifuged for 15 seconds to bring the sample to the bottom of the tube and was then incubated in a 100°C water bath for 5 minutes followed by a centrifugation for five minutes at 13,200 rpm. Lysates were loaded into 10% acrylamide SDS-PAGE gels that were run at a constant current setting of 20 mA per gel at 165V. Gels were transferred to nitrocellulose membrane in a BioRad Trans-Blot Turbo Transfer system. Blots were probed overnight at 4°C in 4% milk in western wash buffer (1x PBS + 250 mM NaCl + 0.1% Tween-20) with mouse monoclonal anti-HA antibody (12CA5, Gift of David Toczyski, University of California, San Francisco), polyclonal anti-Nap1 antibody, polyclonal anti-Ypk1 antibody (Alcaide-Gavilan et al., 2020), or polyclonal rabbit anti-T662P antibody (Gift from Ted Powers, University of California, Davis). Western blots using anti-T662P antibody were first blocked using TBST (10 mM Tris-Cl, pH 7.5, 100 mM NaCl, and 0.1% Tween-20) + 4% milk, followed by one wash with TBST, then overnight incubation with anti-T662P antibody in TBST + 4% BSA. Western blots were incubated in secondary donkey anti-mouse (GE Healthcare NA934V) or donkey anti-rabbit (GE Healthcare NXA931 or Jackson Immunoresearch 711-035-152) antibody conjugated to HRP at room temperature for 60-90 min before imaging with Advansta ECL chemiluminescence reagents in a BioRad ChemiDoc imaging system.

### Log Phase Nutrient Shift

Cells were grown to mid-log phase (OD_600_ 0.4-0.7) in rich carbon (YPD) at 22°C. Cells were shifted to poor carbon (YPG/E) using a vacuum filter apparatus with a 0.45 μM HAWP mixed cellulose ester filter. Cells were washed three times with room temperature YPG/E and then resuspended in room temperature YPG/E by pipetting and grown in a shaking water bath at 25°C.

### Western blot quantification

Western blots were quantified using BioRad Imagelab software v6.0.1. For the elutriation experiments, relative signal was calculated by as a ratio of the signal each time point over the signal at either the zero time point (Whi5) or the ten minute time point (Cln3) (see figure legend for details). The signal was not normalized to the loading control because the loading control signal increased with growth during G1 phase. No Cln3 signal was detected at the zero minute time point in elutriated cells, so the ten minute time point was used as reference for that set of samples. Log phase samples were quantified by setting all signal relative to either the 0 min time point (Figure 2) or wild type cells in rich carbon. The signal was normalized to signal for the loading control band to correct for differences in total protein between samples.

### ^35^S–Methionine pulse labeling

Cells were grown overnight at room temperature to mid-log phase in CSM–methionine media containing 2% dextrose. The culture was concentrated to an OD600 of 0.7 and incubated in a shaking water bath at 25°C. The vehicle or drug was added to the appropriate culture flask. A 1.2 ml sample of culture was transferred to a 1.6 ml screw top tube to label proteins made during the next 15 minutes. The labeling reaction was initiated by adding 1 μl of EasyTag L-[^35^S]-Methionine from PerkinElmer at a stock concentration of 1 μCi/μL. The sample was mixed by vortexing and then placed into the 25°C shaking water bath. Labeling reactions were allowed to progress for 15 min, during which time samples were mixed by inversion every ~5min. Labeling reactions were performed during 15 minute intervals starting every 20 minutes. Samples were centrifuged at 13,200 rpm for 30 sec, the supernatant was removed, and 250 μL of acid-washed beads were added before freezing on liquid nitrogen. Samples were prepared for SDS-PAGE similarly to samples described for western blotting. After running the gel to the dye front, polyacrylamide gels were stained in R-250 Coomassie stain and then dried on Whatman paper using a BioRad Gel Dryer. The dried gel was exposed film.

## Figure Legends

**Figure 1 – figure supplement 1:**
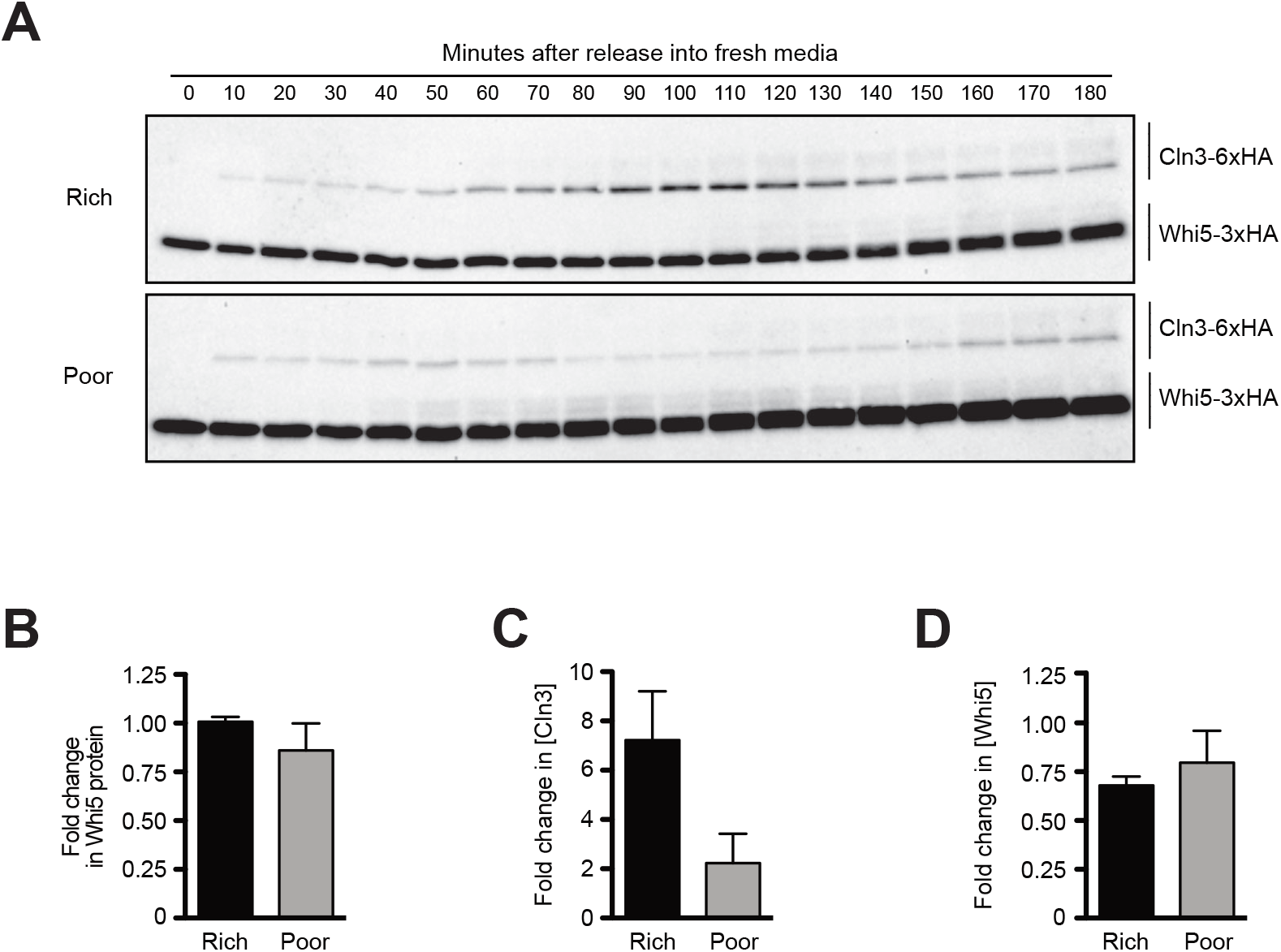
**(A)** A longer exposure of the western blots from **Fig. 1D**. **(B)** The fold change in Whi5 protein levels was calculated by taking the ratio of the Whi5 signal from the time point with peak Cln3 over the Whi5 signal from the zero minute time point. The data show the average of three biological replicates. **(C)** The fold change in Cln3 concentration before bud emergence was calculated by taking the ratio of the value for peak Cln3 concentration over the value from the 10 min time point. The data show the average of three biological replicates. **(D)** The fold change in Whi5 concentration before bud emergence was calculated by taking the ratio of the value for Whi5 concentration at peak Cln3 levels over the value from the zero min time point. The data show the average of three biological replicates. Error bars represent SEM.

**Figure 3 – figure supplement 1:**
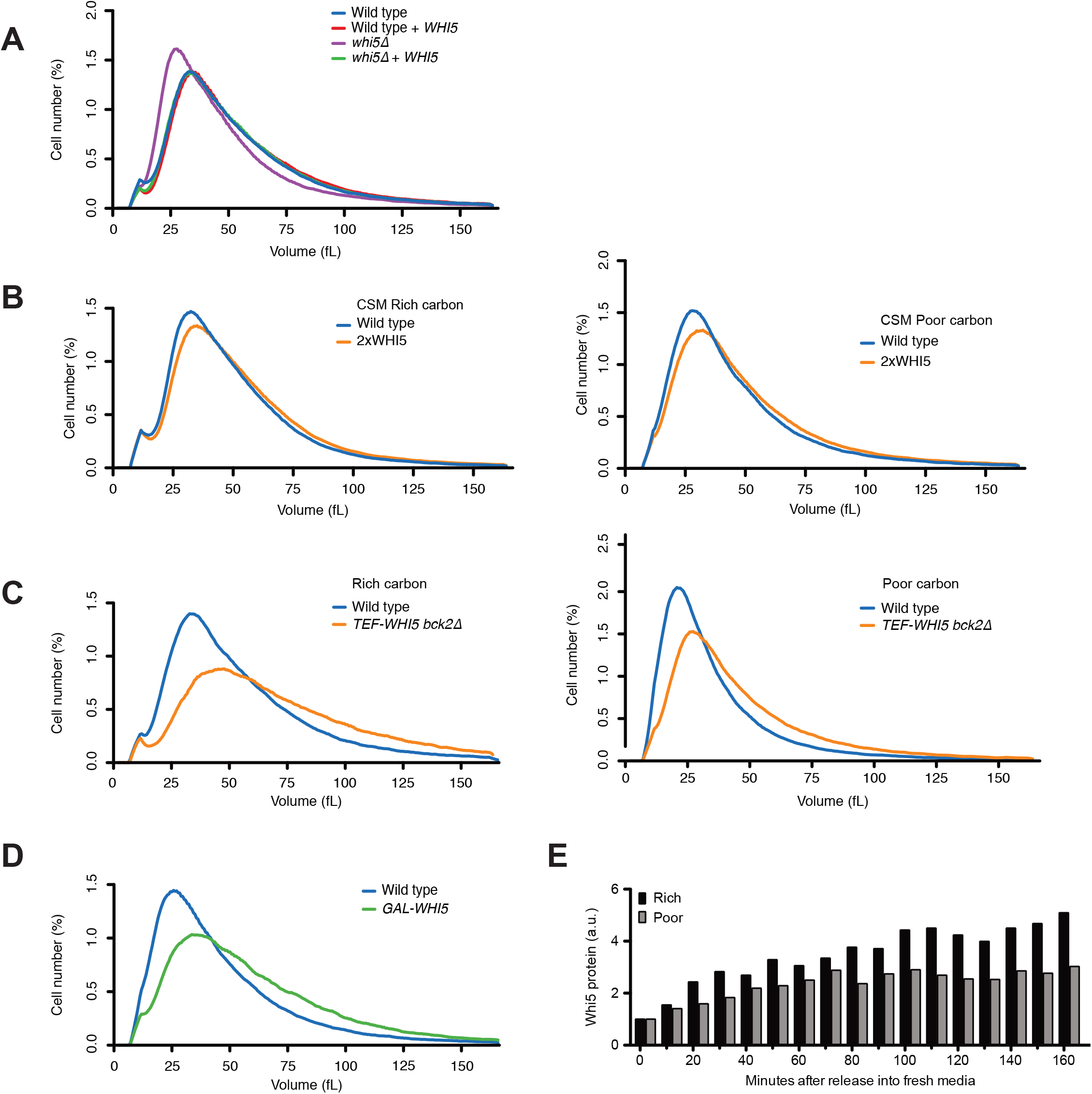
**(A)** Cells of the indicated genotypes were grown to log phase in YPD and cell size was measured with a Z2 Coulter Channelyzer. **(B)** Cells were grown to log phase in complete synthetic media with 2% dextrose (rich carbon) or 2% glycerol and 2% ethanol (poor carbon) and cell size was measured with a Z2 Coulter Channelyzer. **(C)** Cells of the indicated genotypes were grown to log phase in rich (YPD) or poor (YPG/E) carbon and cell size was measured with a Coulter counter. **(D)** Wild type and *GAL-WHI5* cells were grown in YPGal overnight to mid log phase and cell size was measured using a Coulter Counter. **(E)** Whi5 protein levels were quantified from the blots in **Fig. 3D** relative to the 0 min time point in rich or poor.

**Figure 4 – figure supplement 1:**
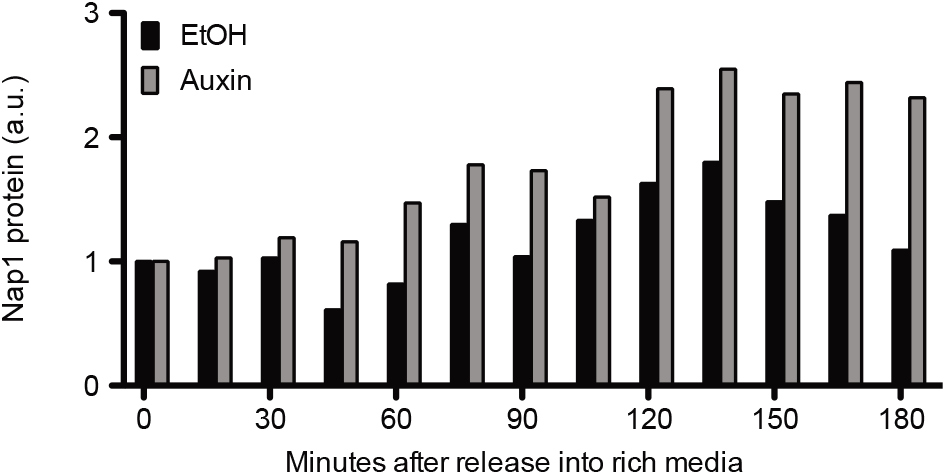
Quantification of Nap1 protein levels in the western blots in **Fig. 4E**. Changes in Nap1 levels were quantified by taking a ratio of the Nap1 signal at each time point over the Nap1 signal at the zero time point.

**Figure 5 – figure supplement 1:**
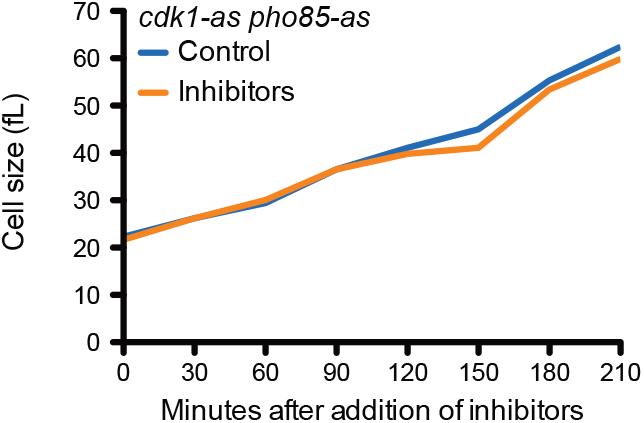
*cdk1-as pho85-as* cells were grown to mid log phase in poor carbon (YPG/E) and small unbudded cells were isolated via centrifugal elutriation. The cells were divided into two cultures and were then released into rich carbon (YPD, not supplemented with additional adenine (see Methods)). 10μM 1-NA-PP1 and 10μM 1-NM-PP1 were added to one culture. Median cell size was measured using a Coulter counter and plotted as a function of time.

**Figure 6 – figure supplement 1:**
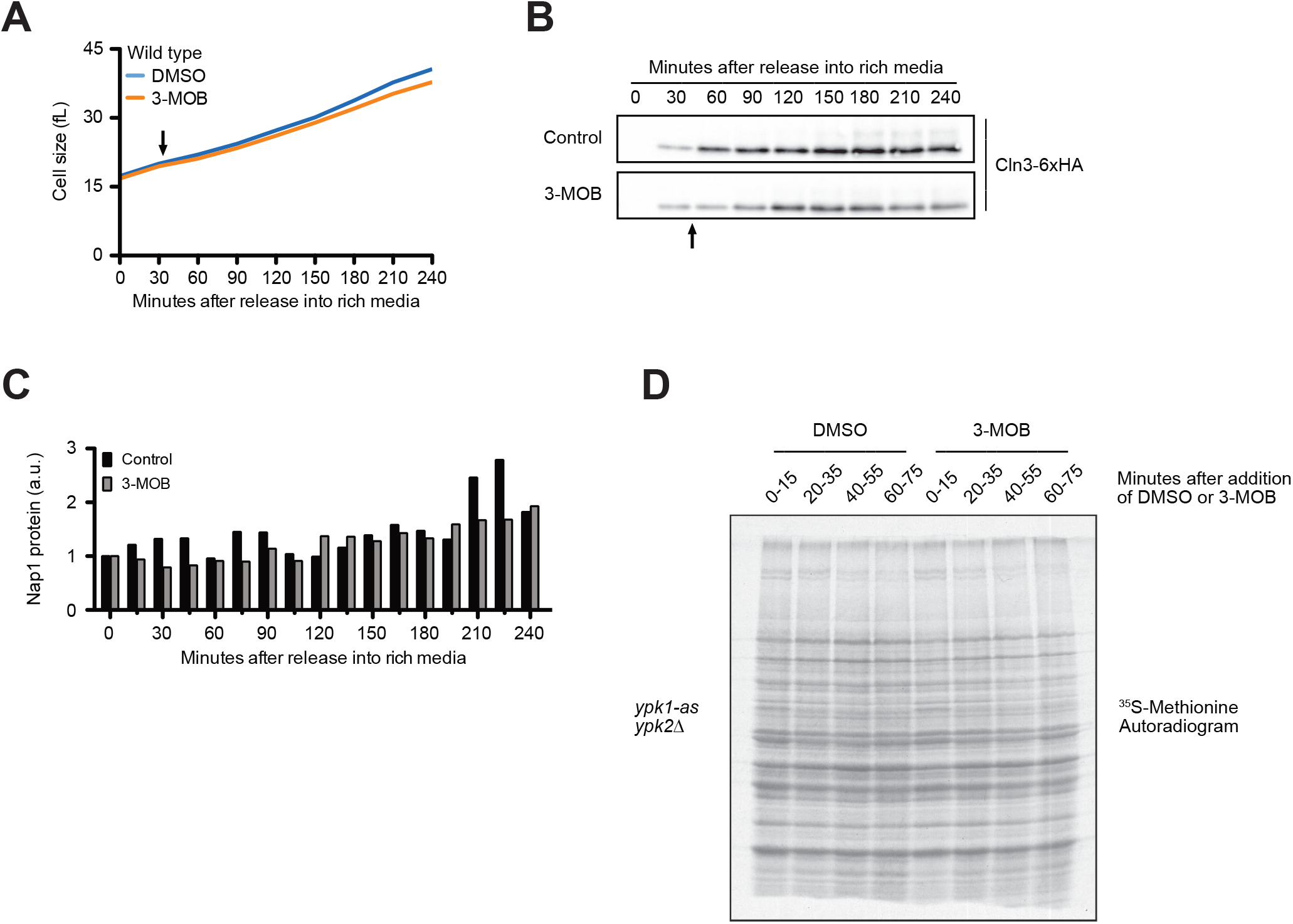
**(A-B)** Wild type cells were grown to mid log phase in poor carbon (YPG/E) and small unbudded daughter cells were isolated by centrifugal elutriation. The cells were split into two cultures and released into rich carbon at 25°C. After 30 min, 25 μM 3-MOB-PP1 was added to one culture (arrow). **(A)** Median cell size was measured using a Coulter counter and plotted as a function of time. **(B)** The behavior of Cln3-6xHA was analyzed by western blot. Arrow indicates when 3-MOB-PP1 was added. **(C)** Quantification of anti-Nap1 loading control blots from **Fig. 6C**. Protein levels at each time point represent a ratio over the signal from the zero minute time point. (**D**) Autoradiogram of ^35^S-methionine labeling to detect *de novo* protein synthesis. *ypk1-as ypk2Δ* cells were grown overnight to mid log phase in-MET synthetic media containing glucose. After addition of 3-MOB-PP1 or vehicle, 1.2 ml samples collected from the cultures were labeled with ^35^S-methionine for 15 min intervals starting every 20 min after addition of 3-MOB-PP1 to measure the rate of protein synthesis within the 15 min intervals.

**Figure 7 – figure supplement 1:**
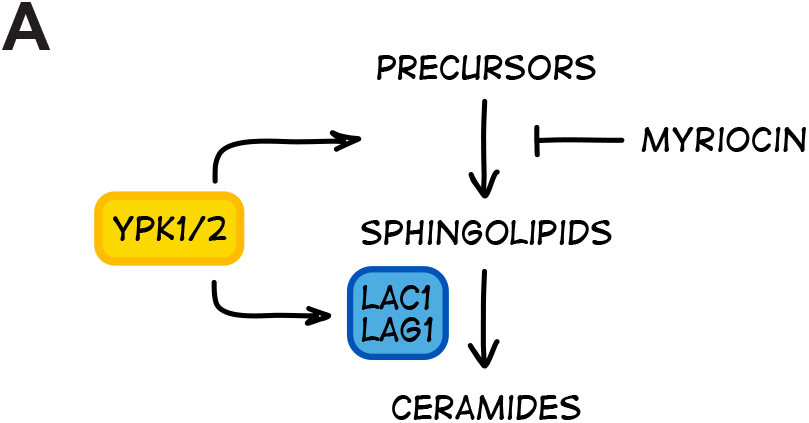
**(A)** A simplified model of the mechanisms by which Ypk1/2 control synthesis of sphingolipids and ceramide.

## Notes

### Competing Interest Statement

The authors have declared no competing interest.

### Summary of Updates

Text edits and addition of several new figures.

